# Selective ultrasonic suppression of pathological brain hyperactivity via an endogenous CAV3–NCX1 axis

**DOI:** 10.64898/2026.11.11.765847

**Authors:** Keunhyung Lee, Jung Moo Lee, Ho-Jeong Kim, Minseok Koo, Seolhee Lee, Sangyep Shin, Wongu Youn, Tien Thuy Phan, C. Justin Lee, Jinhyoung Park, Joo Min Park

## Abstract

Selective suppression of pathological hyperactivity is a major therapeutic challenge across brain disorders. Ultrasound stimulation can modulate neural activity, yet the molecular basis of its inhibitory effects and its ability to selectively inhibit hyperactive rather than basally active cells remain unclear. Here, we identify the caveolin-3 (CAV3)-sodium-calcium exchanger-1 (NCX1) complex as a previously unknown mediator of state-dependent ultrasound-induced neural inhibition. Prolonged ultra-low-intensity ultrasound stimulation engages NCX1-mediated Ca^2+^ extrusion in hyperactive neurons and astrocytes, sparing cells with basal activity. NCX1 is both necessary and sufficient for this inhibitory response, and disruption of the CAV3–NCX1 interface abolishes the ultrasound-induced effects. In vivo, targeted ultrasound stimulation suppresses fear-evoked hyperactivity and freezing in an NCX1-dependent manner and attenuates neuropathic pain-associated hyperactivity, allodynia, and hippocampal epileptiform activity. These findings establish a molecular mechanism for state-dependent ultrasound inhibition, providing a basis for a new approach to neuromodulation that selectively targets pathological hyperactivity.

---

Pathological neural hyperactivity contributes to diverse brain disorders, including epilepsy, chronic pain, and neuropsychiatric conditions^1,2^. Although pharmacological treatments, including antiseizure medications, and neuromodulatory interventions, including transcranial magnetic stimulation, deep brain stimulation, and transcranial electrical stimulation, can reduce excessive neural activity and alleviate symptoms^3–7^, they may also disrupt basal neural activity, limiting therapeutic efficacy or causing adverse effects^8,9^. Thus, selectively suppressing pathological hyperactivity while preserving basal neural activity represents an important therapeutic goal.

Among neuromodulation techniques, transcranial ultrasound has attracted considerable attention for its ability to induce both excitatory and inhibitory neural responses^10–13^. Mechanistic studies have largely focused on excitation, implicating mechanosensitive channels, including Piezo1 and the transient receptor potential channels TRPA1 and TRPC6^14–19^. Ultrasound stimulation can also suppress neural activity and alleviate hyperexcitability-associated responses, including seizures^12^. However, stimulation parameters for inducing inhibition have largely been determined empirically, resulting in substantial variability in inhibitory outcomes across stimulation conditions^12,13^. Although ultrasound can directly activate the mechanosensitive K^+^ channel TRAAK^20^, to our knowledge, the mechanism by which ultrasound selectively suppresses hyperactive cells has not been identified. Achieving reliable neural inhibition requires determining whether ultrasound can selectively suppress hyperactive cells based on their pre-existing state and identifying the molecular machinery linking mechanical stimulation to the suppression of elevated intracellular Ca^2+^ levels.

The sodium-calcium exchanger-1 (NCX1) is a compelling candidate for the ultrasound-induced reductions in elevated intracellular Ca^2+^ levels because it mediates high-capacity Ca^2+^ extrusion driven by an inward Na^+^ gradient and can be modulated by mechanical forces^21,22^. NCX1 localizes within caveolae, where caveolin scaffolding proteins govern its function^23,24^. Although caveolin-1 (CAV1) is abundantly expressed in the brain and proposed as a mechanosensor^25,26^, cardiomyocytes exclusively utilize caveolin-3 (CAV3) to regulate NCX1^23,24^. Whether a caveolin– NCX1 mechanotransduction axis operates in brain cells, and critically, which caveolin isoform directly engages NCX1, remains unknown.

Here, we show that prolonged ultra-low-intensity (<1 W/cm^2^) ultrasound selectively suppresses elevated intracellular Ca^2+^ levels in hyperactive neurons and astrocytes through forward-mode NCX1-mediated Ca^2+^ extrusion. Disrupting caveolin–NCX1 coupling abolishes this response, and structural modelling identifies CAV3, rather than CAV1, as the candidate NCX1-interacting caveolin isoform. In vivo, targeted ultrasound stimulation suppresses fear-evoked neuronal hyperactivity and freezing behavior in an NCX1-dependent manner. It further attenuates neuronal hyperactivity and allodynia in a murine model of neuropathic pain, as well as epileptiform activity in a seizure model. Together, these findings identify the CAV3–NCX1 axis as a molecular mechanism underlying state-dependent ultrasound-induced inhibition of neuronal hyperactivity.

## Results

### Prolonged ultrasound stimulation selectively reduces intracellular Ca^2+^ levels in astrocytes and neurons with elevated basal Ca^2+^ levels

Calcium imaging of cortical astrocytes was performed using Ultrasonocoverslips, with astrocytes cultured directly on the transducer surface (Fig. 1a)^15^. We first established stimulation paradigms that elicited excitatory or inhibitory Ca^2+^ responses in astrocytes (Fig. 1b,c). Excitatory Ca^2+^ responses were evoked with short pulses (6 MHz, 0.15 MPa, I_SPTA_ = 41 mW/cm^2^, 0.5 s), consistent with TRPA1-mediated glutamate release.^15^ By contrast, reducing the pressure to 0.05 MPa and extending the duration to 1–5 s resulted in inhibitory decreases in Ca^2+^ concentrations (Fig. 1b– d). The Ca^2+^ peak amplitude of the inhibitory response increased with burst duration (Fig. 1e), and the fraction of responsive cells rose from 19% at 1 s to 40% at 5 s (Supplementary Fig. 1d). Stimulation was capped at 5 s to maximize inhibitory efficacy while avoiding thermal confounding (Supplementary Fig. 1a,b)^27^. The inhibitory effect depended strongly on baseline Ca^2+^ levels, with greater suppression observed in cells with higher baseline levels (Fig. 1f,g). Responsive cells showed a significant negative correlation between baseline calcium and peak response, in contrast to the weak positive correlation in quiescent cells (Fig. 1h), and had significantly higher baseline calcium levels overall (Fig. 1i). This state-dependent selectivity suggests that prolonged ultrasound stimulation preferentially targets hyperexcitable cell states.

**Fig. 1.**
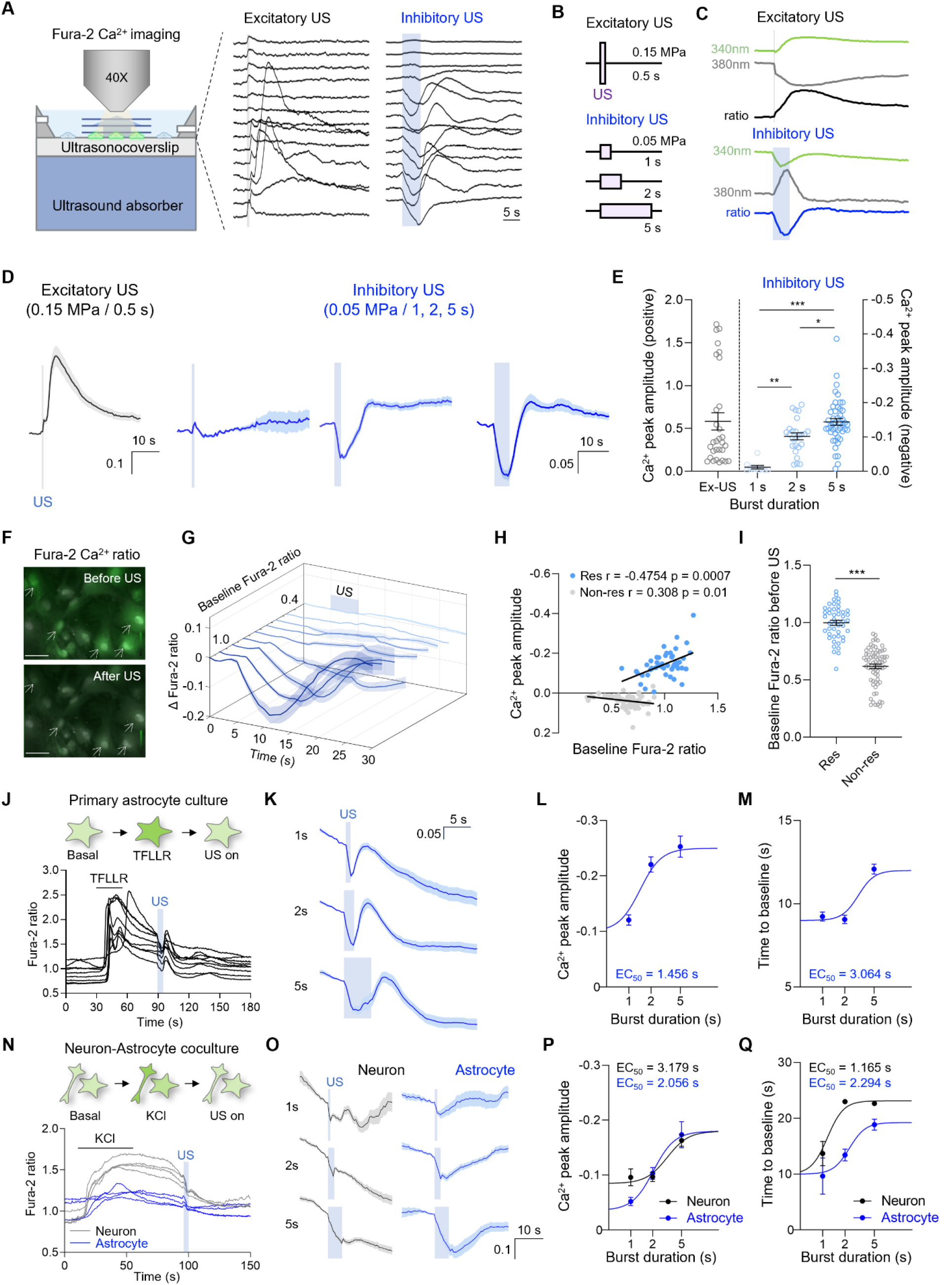
Prolonged low-intensity ultrasound stimulation selectively suppresses intracellular Ca^2+^ concentrations in hyperactive neurons and astrocytes. **a,** Schematic of the in vitro Ca^2+^ imaging setup using the Ultrasonocoverslip and representative calcium responses evoked by application of excitatory or inhibitory ultrasound (US) in a single Ultrasonocoverslip. Scale bar, 5 s. **b,** Acoustic pressure and total burst period determining excitatory or inhibitory US used in the experiment. **c,** Representative Fura-2 AM fluorescence traces at 340 nm (green) and 380 nm (grey) excitation wavelengths and the resulting Fura-2 ratio (F_340_/F_380_) during excitatory ultrasound (top, black) and inhibitory ultrasound (bottom, blue). **d,** Averaged Ca^2+^ traces for excitatory (black) and inhibitory (blue) ultrasound conditions. Blue shading denotes the ultrasound delivery period. Scale bars, excitatory 0.1, inhibitory 0.05, Fura-2 ratio units, 10 s. **e,** Summary of Ca^2+^ peak amplitude for responsive cells under excitatory (left y-axis, grey) and inhibitory (right y-axis, blue) conditions across burst durations. Kruskal–Wallis test with Dunn’s multiple comparison test. **f,** Representative calcium images showing inhibitory ultrasound suppresses calcium responses from hyperactive astrocytes. Arrows indicate astrocytes that react to inhibitory ultrasound stimulation. Scale bar, 50 µm. **g,** Baseline calcium ratio-dependent astrocytic Ca^2+^ responses to inhibitory ultrasound stimulation. Traces show mean ΔFura-2 ratios grouped by their baseline Fura-2 ratio. **h,** Scatter plot of Ca^2+^ peak amplitude versus baseline Fura-2 ratio between responsive (Res) and non-responsive (Non-res) cells during inhibitory ultrasound stimulation. Pearson correlation (Res): r = −0.4754, *p* = 0.0007, n = 47. (Non-res): r = 0.308, *p* = 0.01, n = 75. Black lines indicate linear regression fit. **i,** Summary of baseline Fura-2 ratio before ultrasound onset in responsive (Res, n = 47) and non-responsive (Non-res, n = 75) cells during stimulation. Mann–Whitney unpaired test. **j,** (top) Experimental scheme: Fura-2-loaded astrocytes were sequentially treated with TFLLR (30 µM) to elevate the intracellular Ca^2+^ concentration, followed by inhibitory ultrasound stimulation. (bottom) Representative individual Ca^2+^ traces from TFLLR-treated astrocytes showing the TFLLR-evoked Ca^2+^ rise and the subsequent ultrasound-induced suppression of Ca^2+^ levels. **k,** Averaged Ca^2+^ traces from TFLLR-treated astrocytes during inhibitory ultrasound stimulation for 1 s (top), 2 s (middle), and 5 s (bottom). Scale bars, 0.05 Fura-2 ratio units, 5 s. **l,** Summary of Ca^2+^ peak amplitudes in TFLLR-treated astrocytes across burst durations. The curve is fitted as a Hill equation with an EC_50_ of 1.456 s. **m,** Time to return to baseline in TFLLR-treated astrocytes across burst durations. The curve is fitted as a Hill equation with an EC_50_ of 3.064 s. **n,** (top) Experimental scheme: Fura-2-loaded neuron-astrocyte cocultures were treated with KCl (60 mM) to elevate intracellular Ca^2+^ concentrations, followed by inhibitory ultrasound stimulation. (bottom) Representative Ca^2+^ traces from neurons (black) and astrocytes (blue) during KCl application and ultrasound stimulation. **o,** Averaged Ca^2+^ traces from neurons (black, left) and astrocytes (blue, right) in KCl-treated co-culture during inhibitory ultrasound stimulation for 1 s (top), 2 s (middle), and 5 s (bottom). Scale bars, 0.1 Fura-2 ratio units, 10 s. **p,** Summary of Ca^2+^ peak amplitude comparing neurons (black) and astrocytes (blue) across burst durations. Both curves are fitted as a Hill equation with EC_50_ values of 3.179 s for neurons and 2.056 s for astrocytes. **q,** Time to return to baseline comparing neurons (black) and astrocytes (blue) across burst durations. Both curves are fitted as a Hill equation with EC_50_ values of 1.165 s for neurons and 2.294 s for astrocytes.

To examine the inhibitory response under controlled conditions of elevated baseline Ca^2+^ levels, astrocytic Ca^2+^ levels were raised using the protease-activated receptor 1 (PAR1) agonist TFLLR^28^, and inhibitory ultrasound stimulation was applied during the decay phase (Fig. 1j). Because the ultrasound-induced inhibition depends on pre-existing Ca^2+^ levels (Fig. 1f–i), agonists and depolarization were used to elevate intracellular Ca^2+^ reproducibly, allowing suppression to be quantified relative to the Ca^2+^ excess at ultrasound onset rather than the agonist-evoked peak. Because endogenous Ca^2+^ extrusion occurs during the decay phase, this paradigm allows assessment of whether ultrasound stimulation enhances intracellular Ca^2+^ clearance^1,2^. Responsive cells were defined by the characteristic reciprocal Fura-2 shift confirming a genuine reduction in intracellular Ca^2+^ concentration. Both the fraction of responsive cells and the amplitude of Ca^2+^ reduction increased with sonication duration, with stimulations for 5 s producing a sustained negative Ca^2+^ peak compared to those for 1 or 2 s (Fig. 1k,l and Supplementary Fig. 1e). Baseline Ca^2+^ levels before ultrasound onset were comparable across duration groups, ruling out differential loading as a confounder (Supplementary Fig. 1f). Notably, longer stimulation durations increased the time required for Ca^2+^ levels to return to baseline, indicating a more sustained inhibitory effect (Fig. 1m). Upon ultrasound termination, a transient Ca^2+^ rebound was observed in astrocytes both in the presence and absence of TFLLR (Supplementary Fig. 1g,h).

To determine whether this mechanism applies to neurons, we used astrocyte–neuron co-cultures. Neuronal Ca^2+^ levels were elevated via high-KCl-induced depolarization^29^, after which ultrasound was applied during the decay phase (Fig. 1n). Both neurons and astrocytes showed duration-dependent Ca^2+^ reductions, with 5-s stimulation producing the largest and most sustained responses (Fig. 1o–q), while the fraction of responsive cells also increased with stimulation duration (Supplementary Fig. 1i). In contrast, a transient Ca^2+^ rebound was not observed in neurons following KCl-induced depolarization, despite a clear reduction in intracellular Ca^2+^ concentration during ultrasound stimulation (Supplementary Fig. 1j). The selective presence of this rebound in agonist-stimulated astrocytes, but not in KCl-depolarized neurons, suggests that it reflects the transient unmasking of ongoing agonist-driven Ca^2+^ mobilization upon cessation of ultrasound-driven extrusion, rather than reverse-mode NCX1 activation. Baseline Ca^2+^ levels before ultrasound onset remained consistent across all stimulation durations in astrocytes and neurons, ensuring that the observed effects were not confounded by variations in intracellular calcium levels (Supplementary Fig. 1k). The second-scale kinetics of suppression are inconsistent with rapid ion-channel gating but match the transport kinetics of NCX1^30–36^. Together, these observations implicate a Ca^2+^-extruding transporter rather than an ion channel as the mediator of prolonged ultrasound-induced inhibition.

### Ultrasound-mediated Ca^2+^ reduction in astrocytes is predominantly driven by forward-mode NCX activity

Astrocytic intracellular Ca^2+^ levels are regulated by the high-capacity, mechanosensitive NCX^21,22,37–39^. NCX operates bidirectionally: in the forward mode (Na^+^ in/Ca^2+^ out), it extrudes Ca^2+^, whereas in the reverse mode (Ca^2+^ in/Na^+^ out), the exchanger transports Ca^2+^ into the cell^40,41^. To determine which NCX transport mode mediates the inhibitory response, three selective NCX pharmacological agents were applied to TFLLR-stimulated astrocytes before ultrasound stimulation (Fig. 2a,b): CB-DMB (1–10 µM), a selective forward-mode NCX blocker; SN-6 (30 µM), a selective reverse-mode blocker^42^; and KB-R7943 (30 µM), which at this concentration blocks both modes^43,44^ (Fig. 2b). To validate functional drug loading, we confirmed that forward-mode NCX blockers elevated baseline Ca^2+^ levels after 15 min of loading time, whereas the reverse-mode blocker maintained levels similar to those in the control group (Fig. 2c).

**Fig. 2.**
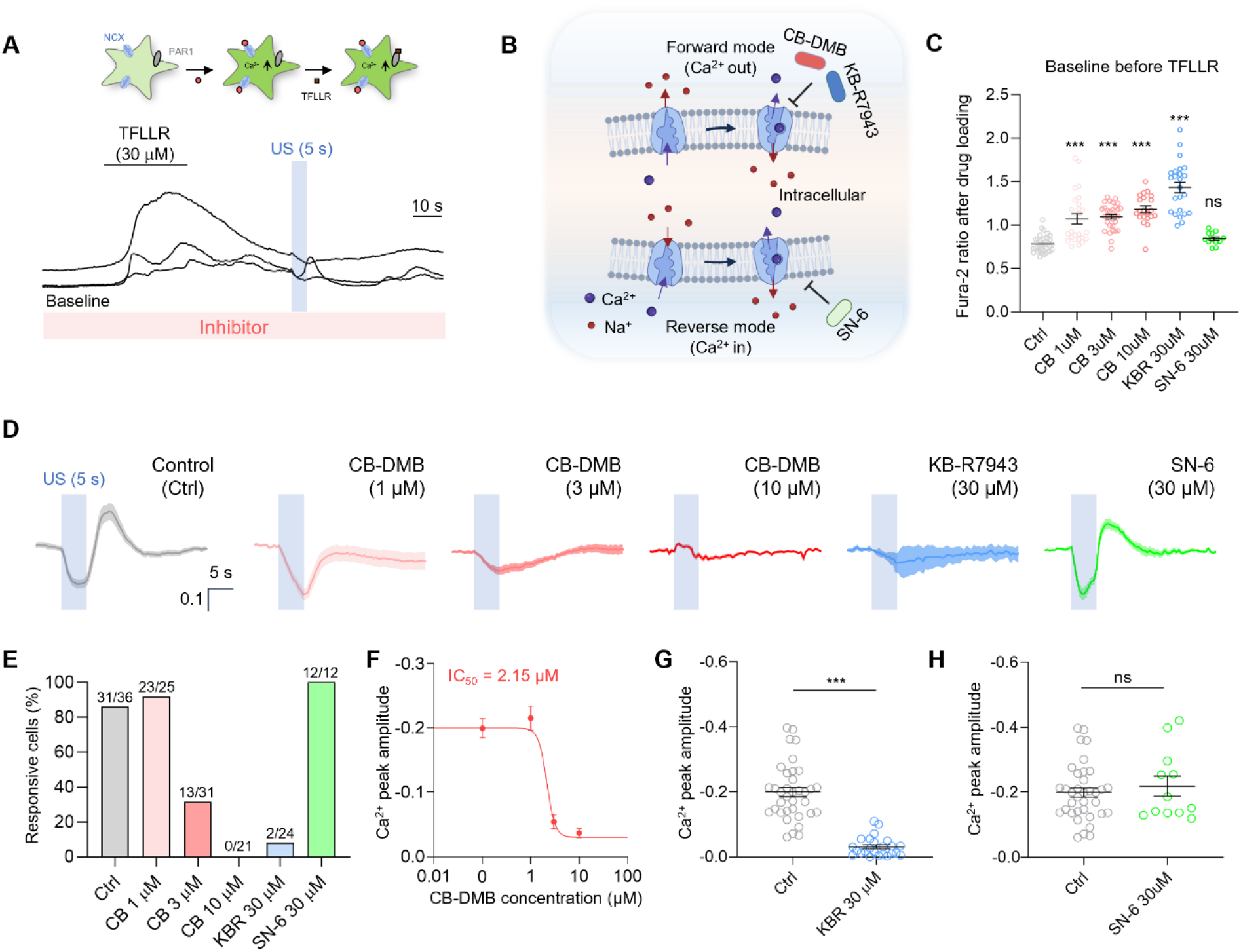
Forward-mode NCX activity mediates ultrasound-induced suppression of intracellular Ca^2+^ levels in astrocytes. **a,** Experimental scheme (top): Fura-2-loaded astrocytes were pre-incubated with a pharmacological NCX inhibitor, and subsequently cells were treated with the PAR1 agonist TFLLR (30 µM) to elevate intracellular Ca^2+^ concentrations, followed by inhibitory ultrasound (US) stimulation for 5 s. Representative Ca^2+^ traces. (bottom): Representative traces showing TFLLR-evoked transient Ca^2+^ increases and the ultrasound-induced response. **b,** Schematic diagram illustrating NCX forward-mode operation (Ca^2+^ out/Na^+^ in, upper) and reverse-mode operation (Ca^2+^ in/Na^+^ out, lower). The binding sites of pharmacological inhibitors are indicated: CB-DMB more selectively blocks forward-mode NCX activity, SN-6 selectively blocks reverse-mode NCX activity, and KB-R7943 (KBR) blocks both forward and reverse modes. **c,** Fura-2 ratio measured after drug loading (baseline before TFLLR application) across all pharmacological conditions: Control (Ctrl, n = 36), CB-DMB 1 µM (n = 25), CB-DMB 3 µM (n = 31), CB-DMB 10 µM (n = 21), KB-R7943 30 µM (n = 24), and SN-6 30 µM (n = 12) conditions. One-way ANOVA with Tukey’s multiple comparison test. **d,** Averaged Ca^2+^ traces from TFLLR-treated astrocytes during inhibitory ultrasound stimulation for 5 s under each pharmacological condition. Blue shading indicates ultrasound delivery period. **e,** Percentage of responsive cells under each pharmacological condition: Ctrl 31/36 (86%), CB-DMB 1 µM 23/25 (92%), CB-DMB 3 µM 13/31 (42%), CB-DMB 10 µM 0/21 (0%), KB-R7943 30 µM 2/24 (8%), and SN-6 30 µM 12/12 (100%). **f,** Dose–response curve of Ca^2+^ peak amplitude versus CB-DMB concentration (0.1–10 µM). The curve is fitted as a Hill equation with an IC_50_ of 2.15 µM. **g,** Summary of Ca^2+^ peak amplitude comparing Ctrl (n = 36) with KB-R7943 30 µM (n = 24). Mann–Whitney unpaired test. **h,** Summary of Ca^2+^ peak amplitude comparing Ctrl (n = 36) with SN-6 30 µM (n = 12). Mann–Whitney unpaired test.

CB-DMB dose-dependently reduced the proportion of responsive cells and Ca^2+^ reduction amplitude (IC_50_ = 2.15 µM; Fig. 2d–f). KB-R7943 similarly abolished the response (Fig. 2d,g), whereas SN-6 had no effect (Fig. 2d,h). Pre-stimulation Ca^2+^ levels were comparable across all conditions, suggesting baseline Ca^2+^ differences were unlikely to account for the observed effects. Additionally, the rebound Ca^2+^ response weas unaffected by SN-6 but suppressed by CB-DMB and KB-R7943 (Supplementary Fig. 2a,b). These results identify forward-mode NCX activity as the predominant mediator of ultrasound-induced Ca^2+^ suppression in astrocytes.

### NCX1 is the primary isoform mediating the ultrasound-induced suppression of Ca^2+^ levels in vitro and in acute brain slices

To definitively establish whether NCX mediates prolonged ultrasound-induced reductions in Ca^2+^ concentrations, we employed genetic ablation of NCX using shRNA. Among the three NCX isoforms differentially expressed across brain regions (Supplementary Fig. 3a–c)^45,46^, western blot analysis identified NCX1 as the predominant isoform in cultured cortical astrocytes (Fig. 3a,b). Isoform-specific knockdown of NCX1 in these cultures (Fig. 3c and Supplementary Fig. 3d–f) altered neither the peak nor the kinetics of TFLLR-induced Ca^2+^ rises (Supplementary Fig. 3h–k). Ultrasound application during the homeostatic decay phase evoked an inhibitory Ca^2+^ response that was reduced by 51% in NCX1-knockdown astrocytes compared to control shRNA-transfected astrocytes (Fig. 3d,e), without affecting pre-stimulation basal Ca^2+^ levels (Fig. 3f).

**Fig. 3.**
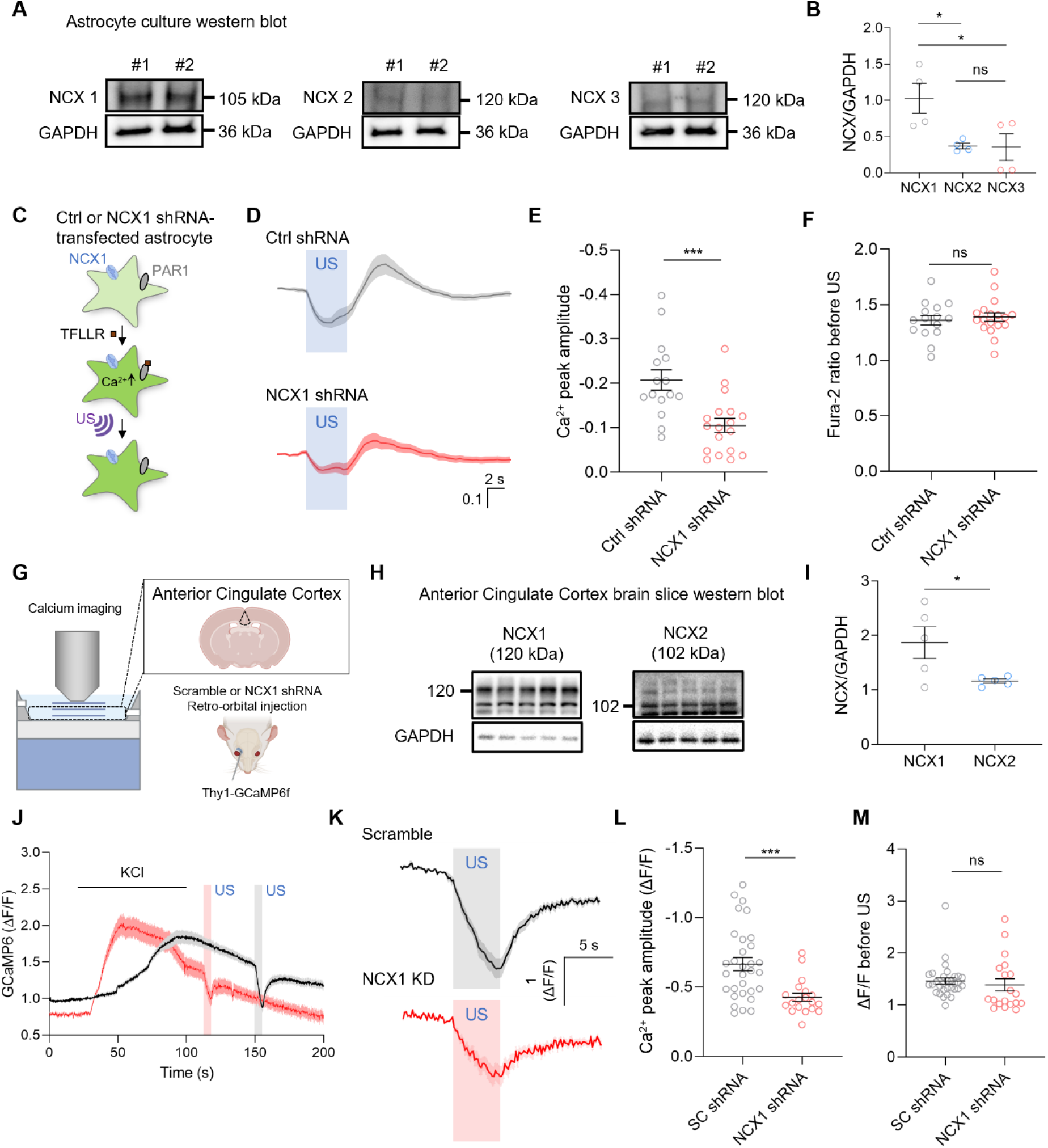
NCX1 is the dominant isoform in cortical astrocytes and ACC neurons, and its genetic ablation impairs the ultrasound-induced reduction of Ca^2+^ concentrations. **a,** Western blots of NCX1 (105 kDa), NCX2 (120 kDa), and NCX3 (120 kDa) protein expression in cultured cortical astrocytes from two independent biological replicates. GAPDH (36 kDa) serves as a loading control. **b,** Quantification of NCX1, NCX2, and NCX3 protein levels normalized to those of GAPDH in cultured cortical astrocytes. n = 4 per group. One-way ANOVA with Tukey’s multiple comparison test. **c,** Experimental scheme: Fura-2-loaded cultured astrocytes transfected with either control (Ctrl) shRNA (scramble) or NCX1 shRNA were activated with TFLLR (30 µM), followed by inhibitory ultrasound (US) stimulation for 5 s. **d,** Averaged Ca^2+^ traces from astrocytes transfected either with Ctrl shRNA (grey, top) or NCX1 shRNA (red, bottom) during 5-s inhibitory ultrasound administration following TFLLR treatment. **e,** Summary of Ca^2+^ peak amplitudes in Ctrl shRNA (n = 15) and NCX1 shRNA (n = 18) astrocytes. Unpaired t-test. **f,** Baseline Fura-2 ratio before ultrasound onset in Ctrl shRNA and NCX1 shRNA astrocytes. Unpaired t-test. **g,** Experimental scheme for acute ACC brain slice imaging of Thy1-GCaMP6f mice. Mice received retro-orbital injection of AAV-PHP.eB encoding scramble or NCX1 shRNA. **h,** Western blot of NCX1 (120 kDa) and NCX2 (102 kDa) protein expression in ACC brain slice tissue. GAPDH serves as a loading control. **i,** Quantification of NCX1 and NCX2 protein levels normalized to those of GAPDH in ACC tissue. NCX1 is the dominant NCX isoform in the ACC. Unpaired t-test. **j,** Time-course of GCaMP6f fluorescence (ΔF/F) in ACC brain slices from Scramble (black) and NCX1 knockdown (red) mice during KCl application and two successive 5-s ultrasound stimulations. **k,** Averaged ΔF/F Ca^2+^ traces from Scramble (black, top) and NCX1 knockdown (red, bottom) ACC neurons during 5-s inhibitory ultrasound stimulation. **l,** Summary of Ca^2+^ peak amplitude (ΔF/F) in ACC slices from SC shRNA (scramble, n = 31) and NCX1 shRNA (n = 19) mice. Mann–Whitney unpaired test. **m,** ΔF/F before ultrasound onset, confirming comparable baseline GCaMP6f signals in both groups. Mann–Whitney unpaired test.

To assess translational relevance, acute brain slices of the anterior cingulate cortex (ACC) from Thy1-GCaMP6f mice were imaged following retro-orbital AAV delivery of scramble or NCX1 shRNA (Fig. 3g). Western blot analysis of mouse ACC confirmed that NCX1 is the prevalent isoform in the ACC, as NCX1 had significantly higher expression levels than NCX2 (Fig. 3h,i). Whole-brain analyses revealed that this NCX1 dominance is consistent across the entire brain (Supplementary Fig. 4a,b). Following KCl-evoked increases in neuronal Ca^2+^ concentration, ultrasound was applied during the decay phase (Fig. 3j). NCX1 knockdown impaired the ultrasound-induced inhibitory Ca^2+^ response, reducing the magnitude of Ca^2+^ suppression by 36% relative to scramble controls (Fig. 3k,l), with comparable pre-stimulation baseline Ca^2+^ levels across groups (Fig. 3m). Collectively, these results identify NCX1 as the primary isoform necessary for ultrasound-induced suppression of Ca^2+^ levels.

### Heterologous NCX1 expression confers ultrasound responsiveness, and intact ion-binding architecture is required

To determine whether NCX1 expression is sufficient to reconstitute the inhibitory Ca^2+^ response, human NCX1A (hNCX1A) was overexpressed in HEK293T cells, which lack detectable endogenous NCX activity under basal conditions^47^. In naïve HEK293T cells, ATP application elevated intracellular Ca^2+^ levels as expected (Fig. 4a). However, the same ATP treatment failed to sufficiently elevate Ca^2+^ concentrations in hNCX1A-overexpressing cells (Supplementary Fig. 5a), likely because tonically active forward-mode NCX1A continuously extruded Ca^2+^ and prevented cytosolic Ca^2+^ accumulation. To circumvent this, intracellular Ca^2+^ concentrations were elevated in hNCX1A-expressing cells by switching to a low-Na^+^ extracellular solution, which reduces the inward Na^+^ gradient driving forward-mode transport and, thereby, promotes reverse-mode Ca^2+^ entry (Fig. 4b). This maneuver selectively raised Ca^2+^ levels in mCherry-positive (hNCX1A-expressing) cells but not in mCherry-negative controls, confirming functional overexpression (Supplementary Fig. 5b). Upon 5-s ultrasound stimulation during the Ca^2+^ decay phase, intracellular Ca^2+^ levels were reduced in naïve HEK293T cells, whereas hNCX1A-overexpressing cells exhibited a significantly greater inhibitory Ca^2+^ response (Fig. 4c–e). Pre-stimulation baseline Ca^2+^ levels were comparable across groups (Supplementary Fig. 5c).

**Fig. 4.**
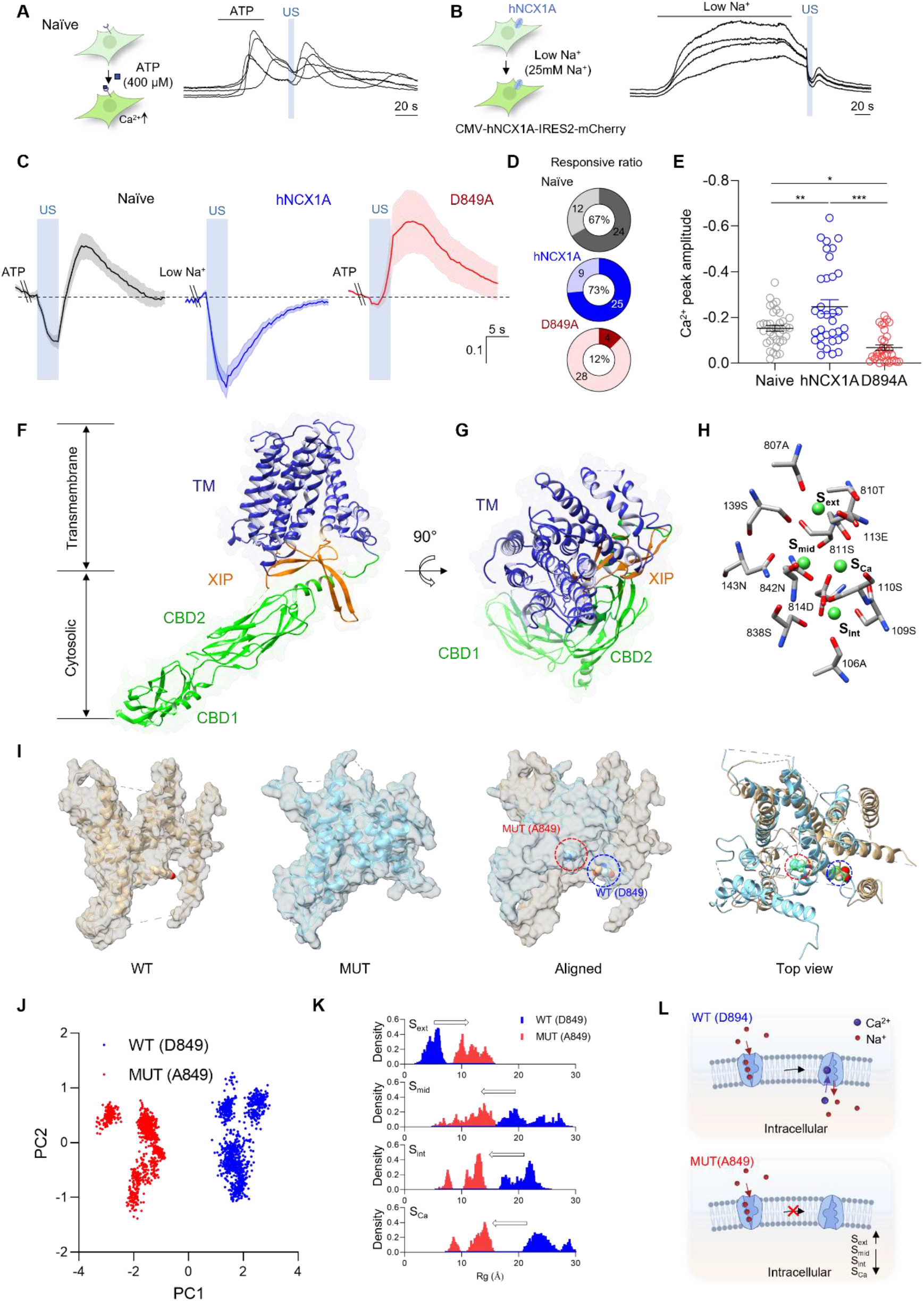
NCX1 expression confers ultrasound responsiveness through an intact ion-binding architecture. **a,** Experimental scheme (left): Naïve HEK293T cells were stimulated with ATP (400 µM) to elevate intracellular Ca^2+^ concentrations. Representative individual Ca^2+^ traces (right) show the ATP-evoked Ca^2+^ rise and the ultrasound-induced Ca^2+^ suppression response. **b,** Experimental scheme (left): HEK293T cells stably expressing human NCX1A (CMV-hNCX1A-IRES2-mCherry) were exposed to low-Na^+^ solution (25 mM Na^+^) to drive Ca^2+^ loading via reverse-mode NCX1 activity, followed by inhibitory ultrasound (US) stimulation for 5 s. Representative individual Ca^2+^ traces (right). **c,** Averaged Ca^2+^ traces during 5-s inhibitory ultrasound stimulation among naïve HEK293T cells (grey), hNCX1A-expressing cells (blue), and D849A mutant-expressing cells (red). **d,** Pie charts showing the percentages of responsive cells: naïve HEK293T (24/36, 67%, grey), hNCX1A-expressing cells (25/34, 73%, blue), and D849A mutant-expressing cells (4/32, 12%, red). Dark wedges, responsive cells; light wedges, non-responsive cells. **e,** Summary of Ca^2+^ peak amplitudes in naïve (n = 36), hNCX1A-expressing HEK293T (n = 34), and D849A mutant (n = 32) cells. One-way ANOVA with Tukey’s multiple comparison. **f,** Three-dimensional structural model of NCX1.3 shown in side view. The transmembrane (TM) domain (blue) spans the lipid bilayer. The inhibitory XIP region (orange) and cytosolic Ca^2+^-binding domains CBD1 and CBD2 (green) are indicated. **g,** Top-down view of the NCX1.3 structural model (90° rotation from f). Structural models generated using AlphaFold2. **h,** Close-up of the NCX1 ion-coordination network within the TM domain. Key residues defining four ion-binding subsites are labelled: S_ext_, S_mid_, S_int_, and S_ca_. Green spheres represent coordinated Ca^2+^ ions. **i,** Surface representations from MD simulations comparing wild-type NCX1 (WT, beige) and the D849A mutant (MUT, light blue), showing four views. **j,** Principal component analysis (PCA) of MD simulation trajectories. WT (blue) and D849A MUT (red) occupy distinct, non-overlapping regions of the PC1–PC2 conformational space. **k,** Radii of gyration (Rg) distributions for four ion-binding subsites from MD simulation trajectories, comparing WT NCX1 (blue) with D849A MUT (red). **l,** Schematic summary of the functional consequence of the D849A mutation. In WT NCX1 (top), Ca^2+^ extrusion is coupled to Na^+^ influx during forward-mode operation. In D849A MUT (bottom), disruption of the ion-binding architecture favors an outward-facing conformation, potentially impairing productive ion-exchange cycling.

To obtain mechanistic insight into how the ion-binding architecture of hNCX1A contributes to ultrasound-induced Ca^2+^ suppression, we performed structural modelling and molecular dynamics (MD) simulations. We analyzed the human NCX1 structure^48^, revealing a canonical alternating-access architecture that comprises a transmembrane domain, regulatory cytosolic Ca^2+^ binding domains (CBD1 and CBD2), and an exchanger inhibitory peptide (XIP) region (Fig. 4f,g). Within the transport core, a close-up view identifies the critical Ca^2+^ regulatory binding sites, highlighting the coordinating residues at the S_ext_, S_mid_, S_int_, and S_Ca_ positions (Fig. 4h). Structural comparisons between wild-type NCX1 (WT) and the D849A mutant (MUT), targeting a critical ion-coordinating residue, demonstrated significant conformational divergence (Fig. 4i). Principal component analysis of the MD trajectories resolved the WT and MUT ensembles into distinct clusters, indicating a fundamental shift in the exchanger’s conformational state (Fig. 4j). Specifically, the D849A substitution perturbed the geometry of the four ion-binding subsites, as evidenced by shifted radii of gyration (R_g_) distributions (Fig. 4k). These results suggest that the D849A substitution alters the exchanger’s conformal dynamics in a manner that may impair productive ion-exchange cycling required for Ca^2+^ efflux (Fig. 4l).

To functionally validate these structural predictions, we explored ultrasound-mediated reductions in Ca^2+^ concentrations in HEK293T cells transfected with the D849A mutant. While cells expressing WT NCX1 exhibited robust responses, the D849A mutant completely failed to reconstitute ultrasound-mediated decreases in Ca^2+^ levels (Fig. 4c–e), without differences in baseline Ca^2+^ loading between groups (Supplementary Fig. 5c). Additionally, hNCX1A overexpression effectively abolished the Ca^2+^ rebound observed in naïve cells, whereas this rebound remained prominent in the D849A mutant (Supplementary Fig. 5d). Together, these functional and structural data highlight the critical role of the NCX1 ion-binding architecture in ultrasound-induced Ca^2+^ suppression. Combined with the selective effects of forward mode specific perturbation with NCX inhibitors (Fig. 2), these findings further support the preferential involvement of forward-mode NCX1 activity in ultrasound-mediated inhibitory Ca^2+^ responses.

### Structural simulations identify CAV3 as the direct NCX1 docking partner, and competitive disruption of the CAV3–NCX1 interface abolishes the ultrasound-mediated suppression of Ca^2+^ levels

Having established NCX1 forward-mode transport as the effector of ultrasound-mediated Ca^2+^ suppression, we next sought to identify the mechanosensory scaffold that couples ultrasound-induced membrane deformation to NCX1 activation. In cardiomyocytes, NCX1 co-immunoprecipitates exclusively with CAV3, not CAV1 or CAV2^23^, and CAV3 organizes NCX1-containing signaling complexes within caveolae^24^. Although CAV1 is more abundantly expressed in brain cells^25^, we performed structural docking simulations to determine which caveolin isoform directly engages NCX1 at the atomic level, and designed cell-penetrating TAT-conjugated peptides to pharmacologically test this interaction (Fig. 5a–d).

**Fig. 5.**
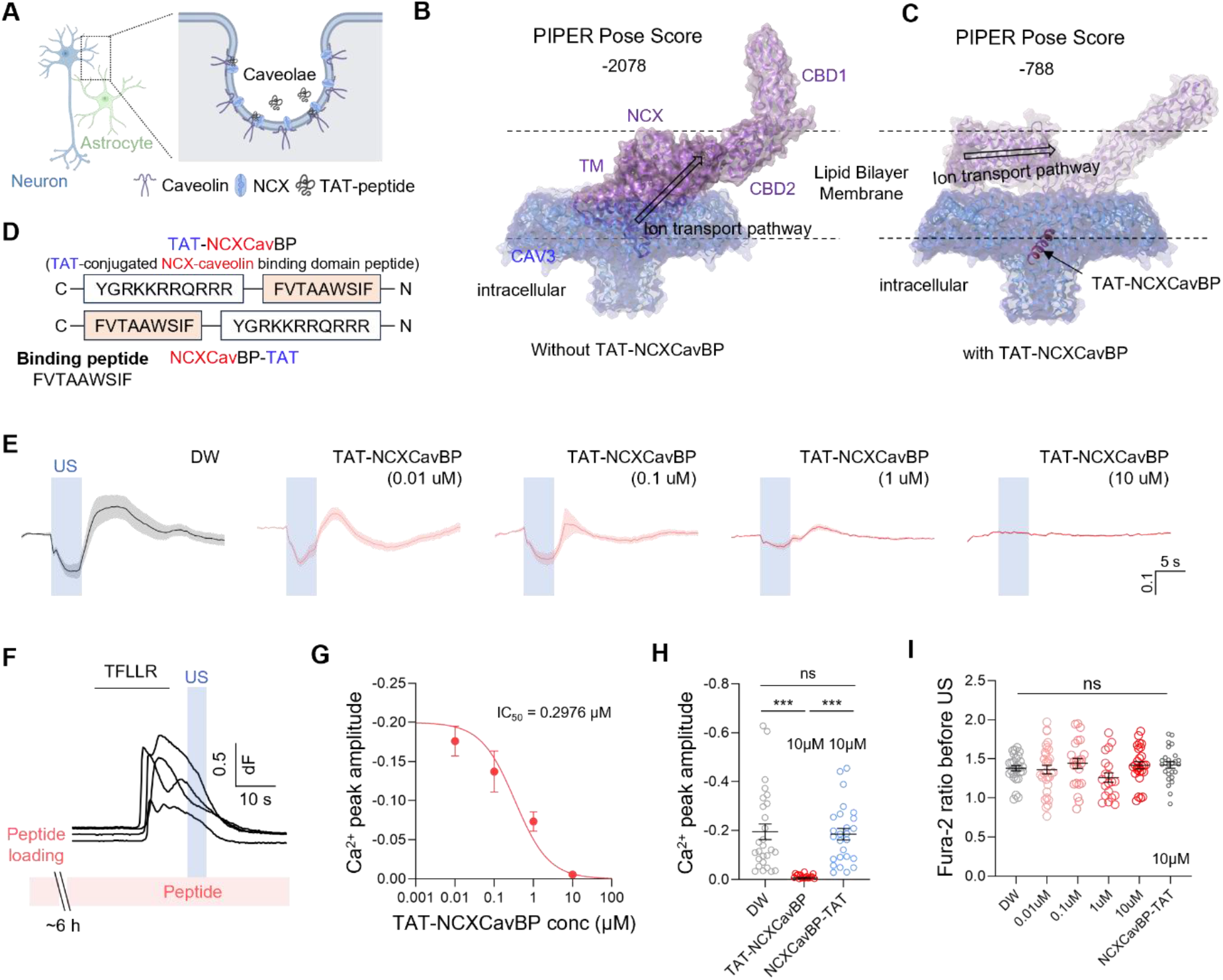
CAV3 is the structural docking partner of NCX1, and competitive disruption of the CAV3–NCX1 interface abolishes the ultrasound-mediated suppression of Ca^2+^ signals. **a,** Schematic of the caveolae membrane microdomain in a neuron–astrocyte pair. CAV3, NCX, and TAT-peptide are indicated. **b,** PIPER rigid-body docking model of NCX1 (purple/blue) and the CAV3 nonamer (blue) in the absence of TAT-NCXCavBP peptide. PIPER pose score = −2,078, indicating energetically highly favorable binding between NCX1 and CAV3. The CBD1, CBD2, and TM domains of NCX1 are labelled. Structural models generated using AlphaFold2. Docking performed using Schrödinger PIPER software. More negative PIPER scores indicate energetically more favorable binding. **c,** PIPER docking model of the CAV3–NCX1 interaction in the presence of TAT-NCXCavBP peptide. PIPER pose score = −788, indicating substantially weaker CAV3–NCX1 binding when TAT-NCXCavBP occupies the NCX1-binding surface on CAV3. Comparison of these two PIPER scores confirms that the competing TAT-NCXCavBP peptide disrupts the CAV3– NCX1 interaction. **d,** Amino acid sequences of the two peptides used in this study: TAT-NCXCavBP (C-YGRKKRRQRRR-FVTAAWSIF-N; TAT-conjugated NCX-caveolin binding domain peptide, active competitor) and NCXCavBP-TAT (C-FVTAAWSIF-YGRKKRRQRRR-N; reversed-orientation control peptide). The CBP core sequence FVTAAWSIF is highlighted. **e,** Averaged Ca^2+^ traces from TFLLR-treated astrocytes during 5-s inhibitory ultrasound (US) stimulation after pretreatment with vehicle (DW, n = 27), TAT-NCXCavBP 0.01 µM (n = 31), TAT-NCXCavBP 0.1 µM (n = 23), TAT-NCXCavBP 1 µM (n = 20), or TAT-NCXCavBP 10 µM (n = 27). **f,** Experimental scheme: Fura-2-loaded astrocytes were preloaded with TAT-NCXCavBP peptide for ∼6 h, then treated with TFLLR (30 µM), followed by inhibitory ultrasound stimulation for 5 s. **g,** Dose–response curve of Ca^2+^ peak amplitude versus TAT-NCXCavBP concentration (0.001–10 µM). The curve is fitted as a Hill equation with an IC_50_ of 0.2976 µM. **h,** Summary of Ca^2+^ peak amplitude comparing vehicle (DW, n = 27), TAT-NCXCavBP (10 µM, n = 27), and NCXCavBP-TAT (10 µM, n = 26) in astrocytes. TAT-NCXCavBP (10 µM) significantly attenuates ultrasound-induced Ca^2+^ suppression, whereas NCXCavBP-TAT has no significant effect. Kruskal–Wallis test with Dunn’s multiple comparison test. **i,** Pre-stimulation Fura-2 ratio before ultrasound onset across all TAT-NCXCavBP concentrations and NCXCavBP-TAT (10 µM), confirming comparable baseline Ca^2+^ levels across all experimental groups. One-way ANOVA.

Rigid-body in silico docking of the NCX1 structure^48^ against an AlphaFold2-modeled CAV3 nonamer^49^ predicted an interaction between CAV3 and NCX1 at the membrane-proximal surface, yielding a robust PIPER pose score of −2,078 (Fig. 5b and Supplementary Fig. 6a–d). To disrupt this interaction, we modeled a competing caveolin-binding peptide (CBP; sequence FVTAAWSIF) derived from the NCX1 caveolin-binding motif (Fig. 5d)^50^. Ternary docking of the CAV3–CBP– NCX1 complex in the presence of TAT-conjugated CBP (TAT-NCXCavBP; YGRKKRRQRRRFVTAAWSIF) yielded a markedly reduced PIPER pose score of −788 (Fig. 5c), structurally confirming that TAT-NCXCavBP competitively destabilizes NCX1 docking onto CAV3. We note that a previous report^51^ raised caveats regarding the specificity of caveolin-binding motif interactions; the atomic-level PIPER docking data presented here provide structural specificity evidence beyond motif-based predictions alone.

To test this prediction experimentally, we applied TAT-NCXCavBP peptide to TFLLR-treated astrocytes at increasing concentrations (0.01–10 µM) and measured the ultrasound-induced suppression of Ca^2+^ levels. While prolonged ultrasound stimulation robustly suppressed Ca^2+^ signals in vehicle-treated cells (distilled water [DW]), TAT-NCXCavBP attenuated this inhibitory response in a dose-dependent manner (IC_50_ = 0.2976 µM), achieving near-complete abolition at 10 µM (Fig. 5e–g and Supplementary Fig. 6e). As a specificity control, the NCXCavBP-TAT peptide (FVTAAWSIFYGRKKRRQRRR), in which the NCX1-derived sequence is positioned at the C-terminus, did not affect Ca^2+^ responses at 10 µM (Fig. 5h). Pre-stimulation Fura-2 ratios were comparable across all conditions, ruling out baseline loading artefacts (Fig. 5i). Together, these structural and pharmacological findings identify the CAV3–NCX1 protein–protein interface as the functional mechanotransduction node through which prolonged ultrasound stimulation activates Ca^2+^ extrusion and establish the NCX1 caveolin-binding motif as a molecular target for modulating this inhibitory pathway.

### Ultrasound stimulation targeting the ACC suppresses pathologically elevated neuronal Ca^2+^ activity across fear and neuropathic pain states via NCX1

Predator-odor exposure is a well-established paradigm for innate fear in rodents^52,53^, involving activation of ACC neurons that are required for this response^52^. To test whether prolonged ultrasound stimulation suppresses this hyperactivity, we performed simultaneous wide-field Ca^2+^ imaging and ultrasound delivery by implanting the optically transparent transducer over the ACC in anaesthetized Thy1-GCaMP6f mice (Fig. 6a–e and Supplementary Fig. 7)^54^. Vaporized mountain lion urine significantly elevated neuronal Ca^2+^ levels in the ACC, a response markedly suppressed by ultrasound stimulation (Fig. 6f,g). Intracisternally administered Bepridil (Fig. 6d), an NCX inhibitor, attenuated this ultrasound-mediated suppression, and c-Fos immunostaining corroborated that the inhibitory effect depends on in vivo NCX activity (Fig. 6f–i).

**Fig. 6.**
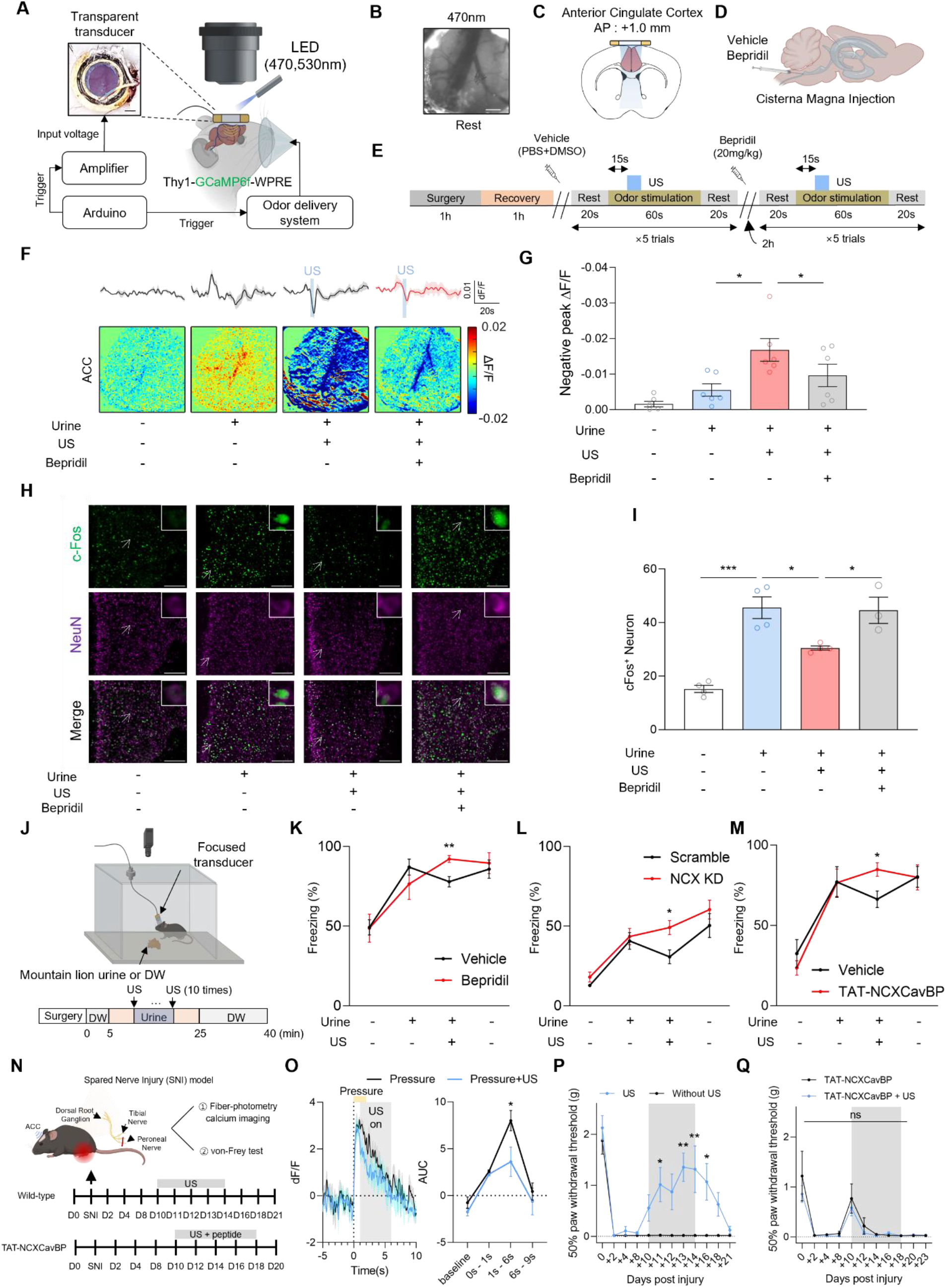
Focused ultrasound stimulation targeting the ACC suppresses pathologically elevated neuronal Ca^2+^ activity across fear and neuropathic pain states in an NCX1-dependent manner. **a,** Experimental schematic for wide-field Ca^2+^ imaging in head-fixed Thy1-GCaMP6f-WPRE mice. A transparent ultrasound transducer is positioned over the ACC cranial window. **b,** Wide-field fluorescence image of the ACC at rest under 470 nm LED illumination. Scale bar, 0.5 mm. **c,** Coronal brain atlas section showing the ACC at AP +1.0 mm from bregma, indicating the ultrasound-targeting location. **d,** Schematic of cisterna magna injection for intrathecal pharmacological delivery of vehicle (PBS+DMSO) or bepridil (20 mg/kg). **e,** Experimental timeline for the wide-field imaging paradigm. Each trial: Rest (20 s) → Odor stimulation (60 s, with 5-s ultrasound (US) stimulation applied 15 s after odor onset) → Rest (20 s). **f,** Wide-field Ca^2+^ imaging pseudo-color ΔF/F maps of the ACC under four conditions: Urine−/US−/Bepridil−, Urine+/US−/Bepridil−, Urine+/US+/Bepridil−, and Urine+/US+/Bepridil+, with averaged ΔF/F time-course traces for each condition. **g,** Summary of negative peak ΔF/F across four conditions. n = 6. One-way ANOVA with Sidak’s multiple comparison test. **h,** Confocal immunofluorescence images of ACC coronal sections stained for c-Fos (green), NeuN (violet), and their merged image under four experimental conditions. Scale bars, 200 µm. **i,** Quantification of c-Fos-positive neurons in the ACC across four conditions. n = 4 (Urine−/US−/Bepridil−), n = 4 (Urine+/US−/Bepridil−), n = 4 (Urine+/US+/Bepridil−), and n =3 (Urine+/US+/Bepridil+). One-way ANOVA with Holm–Sidak’s multiple comparison test. **j,** Schematic of the behavioral fear suppression setup for freely moving mice. A miniaturized wearable focused transducer (5 mm diameter) was affixed to the skull over the ACC. Behavioral timeline: Vehicle (DW) baseline (0–5 min) → Urine exposure (5–25 min, US applied 10 times at 15 s after odor onset) → DW recovery (25–40 min). **k,** Freezing percentage comparing vehicle-treated (black) and bepridil-treated (red) mice across behavioral conditions. Unpaired t-test. n = 7. **l,** Freezing percentage comparing Scramble shRNA (black) and NCX1 knockdown (NCX KD, red) mice across behavioral conditions. Mann–Whitney U test. n = 9. **m,** Freezing percentage comparing vehicle-treated (black) and TAT-NCXCavBP-treated (red) mice across behavioral conditions. Unpaired t-test. n = 6. **n,** Schematic of the spared nerve injury (SNI) model and the experimental timeline. Neuropathic pain states were evaluated via fiber-photometry calcium imaging in Thy1-GCaMP6f mice and von-Frey tests in both wild-type and TAT-NCXCavBP-treated mice. **o,** (left) Fiber photometry Ca^2+^ traces (dF/F) in the ACC comparing pressure stimulation alone (Pressure, black) and pressure stimulation combined with 5-s ultrasound stimulation (Pressure+US, blue). Pressure onset at 0 s; ultrasound active during 1–6 s (grey shading). n = 4. (right) Area under the curve (AUC) analysis of GCaMP6f responses in the ACC across four time windows: baseline, 0–1 s, 1–6 s, and 6–9 s after stimulus onset. Two-way ANOVA with Sidak’s multiple comparison test. n = 4. **p**, 50% paw withdrawal threshold measured via the von-Frey test, in the US-treated (blue, n = 6) and non-US control (black, n = 4) groups. Two-way ANOVA with Sidak’s multiple comparison test. **q,** Comparison of the TAT-NCXCavBP-treated and TAT-NCXCavBP combined with ultrasound stimulation groups across the 23-day postoperative period. Two-way ANOVA with Sidak’s multiple comparison test. n = 3 per group.

To assess whether this suppression of Ca^2+^ signals in the ACC translates into behavioral modulation, we next examined freely moving mice using a miniaturized focused ACC-targeting transducer affixed to the intact skull (Fig. 6j and Supplementary Figs. 8 and 9a). Ultrasound stimulation reduced predator urine-induced freezing behavior in vehicle-treated mice, but this effect was blocked by bepridil (Fig. 6k and Supplementary Fig. 9b). Moreover, ultrasound stimulation alone did not elicit freezing (Supplementary Fig. 9c), confirming that the behavioral effect reflects genuine suppression of fear-evoked activity rather than a nonspecific motor response.

To complement pharmacological inhibition with genetic specificity, NCX1 was selectively knocked down in neuronal cells via retro-orbital injection of AAV-PHP.eB encoding an shRNA under the pSicoR system. Viral expression and knockdown efficiency were confirmed by fluorescence imaging and western blot analyses (Supplementary Fig. 9d–i). Ultrasound stimulation significantly reduced predator-odor-evoked freezing in scramble-shRNA control mice but not in NCX1 knockdown animals (Fig. 6l and Supplementary Fig. 9j). In both groups, fear responses re-emerged after cessation of ultrasound stimulation, confirming that the behavioral rescue depends on active NCX1-mediated Ca^2+^ suppression during stimulation rather than a lasting change in circuit excitability. Furthermore, intracisternally delivered TAT-NCXCavBP, which disrupts CAV3– NCX1 interactions (Supplementary Fig. 9l)^50^, impaired the ultrasound-mediated behavioral rescue (Fig. 6m and Supplementary Fig. 9k), confirming that CAV3-mediated NCX1 localization is required for this therapeutic effect.

To determine whether this inhibitory framework generalizes to alternative states of pathological cortical hyperexcitability, we extended our investigation to a spared nerve injury (SNI) model of neuropathic pain (Fig. 6n and Supplementary Fig. 10c). Fiber photometry calcium imaging confirmed that ultrasound applied at the peak of pain-evoked responses significantly reduced aberrant ACC neuronal Ca^2+^ activity (Fig. 6o), with similar suppression observed in the somatosensory cortex (Supplementary Fig. 10a,b). Prolonged ultrasound stimulation of the ACC reversed mechanical allodynia in WT mice, as assessed by the von-Frey test (Fig. 6p). However, this effect was abolished following TAT-NCXCavBP administration, indicating that the CAV3– NCX1 axis is essential for ultrasound-mediated attenuation of mechanical allodynia (Fig. 6q). Additionally, we tested an acute seizure model induced by intrahippocampal kainic acid (KA) microinjection (Supplementary Fig. 10d,e) along with ultrasound stimulation. Contralateral electrocorticography (ECoG) recordings revealed that localized ultrasound administration at the injection site effectively suppresses the propagation of epileptiform activity, profoundly reducing epileptic spikes (Supplementary Fig. 10f–h).

This suppression closely mirrored that observed in the fear paradigm, suggesting that the NCX1 forward-mode mechanism is not circuit-specific but is broadly recruited whenever ACC neurons are in a state of pathological hyperactivation, regardless of the driving etiology. Collectively, our findings in the innate fear paradigm delineate the molecular mechanisms underlying the CAV3– NCX1 mechanotransduction axis, while extending this approach to neuropathic pain and epileptic seizure models demonstrates its therapeutic versatility. Furthermore, the suppression of hyperactivity confirmed by calcium imaging and electrophysiology establishes region-targeted prolonged ultrasound stimulation as a broadly applicable, cross-modal, and noninvasive strategy for inhibitory neuromodulation across distinct disease contexts.

## Discussion

This study demonstrates that ultra-low-intensity, prolonged ultrasound stimulation (0.05 MPa, 1– 5 s at 6 MHz) reliably suppresses intracellular Ca^2+^ signals in hyperactive astrocytes and neurons via an NCX1-dependent, state-dependent mechanism. By identifying the CAV3–NCX1 caveolar complex as the essential mechanotransduction node, our findings bridge molecular-level biophysics with cross-modal in vivo efficacy. The slow, seconds-range kinetics of this suppression align with the conformational rate-limiting steps of ion transporters rather than rapid channel gating, expanding the known mechanosensitivity of NCX1 to acoustic strain in both neural and glial networks, supported by the abolition of ultrasound-induced responses via CB-DMB treatment, NCX1 knockdown, and the D849A transport-deficient mutant. Because the in vitro experiments used agonists, depolarization, or low-Na^+^ solution to elevate Ca^2+^ experimentally, the physiological and pathological relevance of the modest suppression should be assessed in the light of the in vivo reductions in odor-, injury-, and seizure-evoked activity (Fig. 6 and Supplementary Figs. 9 and 10).

Our use of 1–5-s stimulation contrasts with the brief (<1 s) pulses that activate mechanosensitive ion channels. The 1–5-s window was not prescribed in advance; it emerged from a systematic survey of pressure and duration in which short, higher-pressure pulses evoked excitation and longer, lower-pressure exposures produced suppression (Fig. 1b–e and Supplementary Fig. 1d). The channels implicated in ultrasound-evoked excitation activate within milliseconds and then deactivate: Piezo1 currents inactivate with a fast time constant of ∼12–15 ms at −80 mV^55^, TRPA1 undergoes Ca^2+^-dependent desensitization with a half-time of ∼5 s at physiological extracellular Ca^2+^ concentrations^56^, and TRPC6 is subject to Ca^2+^/calmodulin-dependent inactivation after receptor activation^57^. Channel-mediated Ca^2+^ entry is, therefore, largest at stimulus onset and already waning over the interval in which we observe suppression. As a secondary active transporter, NCX1 exchanges substantially fewer ions per cycle than ion channels conduct, so sustained stimulation is required for cumulative forward-mode Ca^2+^ extrusion to produce a measurable effect, consistent with the progressive increase in both responsive-cell fraction and suppression amplitude from 1-s to 5-s stimulation (Fig. 1k,l). Measured turnover is 1,000–5,000 per s, i.e., a factor of ∼10³ lower than that of single channels^36^, and caveolae flatten passively under membrane tension without ATP or actin^58^, placing the whole CAV3–NCX1 node in the same slow regime. Among the effectors able to lower cytosolic Ca^2+^, NCX1 is the one whose low-affinity, high-capacity forward-mode transport matches both this time domain and the observed state dependence. The 5-s upper bound was set to remain below the thermal threshold (Supplementary Fig. 1a,b)^27^.

A central paradox addressed here is the functional requirement of CAV3, despite CAV1 being the more abundant isoform in the central nervous system^25,26^. Our atomic-resolution docking simulations and in vivo co-immunoprecipitation data resolve this by demonstrating a stereospecific, direct binding interface between NCX1 and CAV3, which provides structural evidence beyond motif-based prediction alone. Crucially, TAT-NCXCavBP competitively destabilizes this binary interface rather than nonspecifically disrupting CAV1 scaffolds. While our co-immunoprecipitation data demonstrate that NCX1 can associate with both CAV1 and CAV3 in brain tissue, the complete loss of ultrasound-mediated suppression upon TAT-NCXCavBP application underscores the CAV3–NCX1 interface as the indispensable functional node for acoustic mechanotransduction.

The competitive displacement mechanism of TAT-NCXCavBP provides important translational implications beyond mechanistic understanding. The NCX1-derived caveolin-binding motif effectively acts as a “molecular key” that occupies the CAV3 docking surface, functioning as a dominant-negative inhibitor to uncouple endogenous Ca^2+^ extrusion from acoustic strain.

The present study focuses on the CAV3–NCX1 axis in cortical astrocytes and ACC neurons, where NCX1 is the dominant isoform. Complementary co-immunoprecipitation experiments now demonstrate that NCX1 associates with both CAV1 and CAV3 in brain tissue. Although our atomic-level docking simulations specifically resolved a direct CAV3–NCX1 interface, the convergence of co-immunoprecipitation, computational, and TAT-NCXCavBP peptide-displacement data establishes that NCX1 engages caveolin scaffolds more broadly, and that disrupting this caveolin–NCX1 interface is sufficient to abolish ultrasound-mediated Ca^2+^ suppression. The generalization of ultrasound-mediated NCX1 activation from fear-related ACC circuits to allodynia-associated cortical hyperactivation and acute epileptogenic network states suggests that the caveolin–NCX1 axis is broadly engaged whenever neurons are in a state of sustained activation, irrespective of the upstream driver. In innate fear, neuropathic pain, and acute seizure models, focused ultrasound stimulation engages the NCX1 efflux machinery state-dependently, sparing quiescent cells and minimizing the risk of over-suppression in normally functioning circuits.

One limitation concerns the 6 MHz carrier used throughout. At this frequency, the focus is in the sub-millimeter range, which allows a single cortical region to be addressed in the mouse, and loss across the thin murine skull is modest. Attenuation in bone rises with frequency^59^, so the same carrier would not usefully penetrate the human skull, and human transcranial ultrasound transducers accordingly operate between approximately 250 and 1,000 kHz^60^. Thus, for clinical translation, the mechanical exposure presented in this study must be preserved at a carrier frequency in this range. Here, we refer to the guidelines provided by the International Ultrasound Neuromodulation Consortium (ITRUSST) as well as the database of acoustic tissue properties provided by the Foundation for Research on Information Technologies in Society (IT‘IS; Zurich, Switzerland). The critical frequency-dependent mechanical considerations are the mechanical index (MI), acoustic radiation force (ARF), and beam size^61^. Since beam size is directly related to the transducer dimensions and target brain region, it is not a relevant factor in this discussion^62^. For the MI calculation, the 0.05 MPa peak negative pressure used here corresponds to an MI of 0.02 at 6 MHz and 0.07 at 500 kHz, both far below the MI ≤ 1.9 regarded as safe for human exposure^61^; the pressure this mechanism requires is, therefore, achievable with a sub-MHz transducer at a wide margin. Similarly, for the ARF calculation, conversion to a 500 kHz carrier frequency from a 6 MHz beam drastically reduces the nominal ARF by a factor of 1.2 to 6.7 at an exponential power of approximately 1.1, depending on the target brain region (IT‘IS Foundation)^63^. This level of reduction is practically easy to recover by increasing the driving voltage of the ultrasound transducer by the appropriate ratio. Nonetheless, whether the suppression described here transfers to such a carrier remains to be tested directly.

The breadth of disease contexts in which hyperactivation plays a role, including chronic pain, anxiety disorders, post-traumatic stress disorder, and certain forms of epilepsy, positions the CAV3–NCX1 axis as a molecular target of considerable therapeutic interest. Noninvasive, transcranial focused ultrasound stimulation can reach the brain cortex through the intact skull with the millimeter-scale precision demonstrated here, and the miniaturized wearable transducer design we describe is compatible with freely moving, awake animals, bringing device translation within reach. Future work should determine whether repeated ultrasound sessions produce lasting reductions in allodynia or anxiety-like behavior through long-term modulation of NCX1 expression or caveolar organization, and whether the approach scales to larger brain volumes in non-human primates. The CAV3–NCX1 axis, thus, serves as the central molecular bridge through which ultrasound drives state-dependent cortical inhibition (Supplementary Fig. 11). The ability to selectively suppress pathological Ca^2+^ signaling in hyperactive brain regions without pharmacological systemic exposure or surgical implants represents a paradigm shift in inhibitory neuromodulation for conditions as diverse as neuropathic pain, innate fear dysregulation, and cortical hyperexcitability disorders.

## Methods

### In vitro stimulation and calcium imaging

Throughout in vitro experiments, an Ultrasonocoverslip was utilized as a cell stimulator^15^. To minimize acoustic reflections, an ultrasound absorber (Aptflex F28, Precision Acoustics) was placed underneath the transducer, which was secured in a custom-made jig enabling buffer circulation and ultrasound transmission simultaneously. Intracellular Ca^2+^ levels were monitored via Fura-2 ratiometric imaging to account for potential cell displacement artifacts. For Fura-2 loading in astrocytes, astrocyte–neuron co-cultures, and HEK293T cells, cells were incubated for 30 min at room temperature in a mixture of Fura-2 AM (F1221, Invitrogen), 20% pluronic acid, and 3 ml of HEPES buffer. HEPES buffer contained 150 mM NaCl, 10 mM HEPES, 3 mM KCl, 2 mM CaCl₂, 2 mM MgCl₂, and 5.5 mM glucose (pH 7.3, 320 mOsmol/kg). To record calcium responses, epi-fluorescence microscopy (BX51W1, Olympus) with a 40× objective lens (LUMPlanFLN 40X, Olympus) and a 2-wavelength LED controller (pE-340fura, CoolLED) was used. The frame rate of image acquisition was 2 Hz. Calcium concentrations were determined by the ratio of fluorescence intensities (F_340nm_/F_380nm_).

### Ultrasound stimulation parameters

Ultrasound stimulation pulses were triggered by the first function generator (33210A, Agilent) sensing the trigger pulse signal generated by a programmable trigger control system (Master-8, A.M.P.I), delivering a square wave having a 1.5-kHz pulse repetition frequency with designated burst duration into the second function generator (33600A, Agilent). After receiving this pulse train, the second function generator transmitted a 6-MHz burst with a 50% duty cycle. This transmission total pulse was delivered into an RF amplifier (75A250A, Amplifier Research) to amplify the electric voltage, which was ultimately transduced into an ultrasound stimulation pulse by the Ultrasonocoverslip with an acoustic negative pressure of 0.05 MPa, corresponding to a spatial peak temporal average intensity of 41 mW/cm².

### Primary astrocyte cultures

Primary astrocytes were isolated from the cerebral cortex of C57BL/6J mouse pups (P0–P1) regardless of sex. The cerebral cortex was dissected in Hank’s Balanced Salt Solution (HBSS, LB 003-02, Welgene), cleared of meninges, and dissociated into a single-cell suspension. Cells were cultured in Dulbecco‘s Modified Eagle Medium (DMEM, 10-013, Corning) supplemented with 25 mM glucose, 4 mM L-glutamine, 1 mM sodium pyruvate, 10% heat-inactivated horse serum (26050-088, Gibco), 10% heat-inactivated fetal bovine serum (FBS; 10082-147, Gibco), and 100 units/ml penicillin–streptomycin (15140-122, Gibco) at 37°C with 5% CO₂. After 3 days in vitro (DIV), the medium was thoroughly washed with repeated pipetting to remove cell debris and non-adherent cells, followed by medium replacement. Astrocytes cultured for 7–14 DIV were detached from the culture plates by trypsinization 1–3 days prior to the experiment. Ultrasonocoverslips were pre-coated with 0.1 mg/ml poly-D-lysine (P6407, Sigma-Aldrich) for 1 day to enhance cell adhesion to the transducer. The detached astrocytes were then reseeded onto the Ultrasonocoverslips and incubated for 1–3 days.

### Primary astrocyte–neuron co-cultures

Primary cortical neurons were isolated from embryos (E14–E16) obtained from pregnant C57BL/6J mice. Following decapitation, the embryonic cortices were dissected and washed three times with HBSS. The tissue was then treated with 0.1% trypsin (15090-046, Gibco) and incubated for 5 min at 37°C. After incubation, heat-inactivated FBS was added to the cortex, and the cortices were gently dissociated by trituration. The resulting cell suspension was centrifuged at 1,000 rpm for 3 min, resuspended in Neurobasal medium (10888022, Gibco) supplemented with 5% heat-inactivated FBS, 2% B27 supplement (17504-044, Gibco), 2 mM GlutaMAX-I (35050-061, ThermoFisher), and 10 units/ml penicillin–streptomycin, and filtered through a 40-µm cell strainer. Neurons (1 × 10⁵ cells) and astrocytes (5 × 10⁴ cells) were then co-cultured on Ultrasonocoverslips pre-coated with 0.1 mg/ml poly-D-lysine (P6407, Sigma-Aldrich). The cells were maintained in culture medium and incubated at 37°C in a humidified 5% CO₂ incubator for 7–10 days prior to the experiment. Half of the medium was refreshed every 2–3 days.

### HEK293T cell cultures

Human embryonic kidney 293T (HEK293T) cells were purchased from ATCC and tested negative for mycoplasma contamination. HEK293T cells were cultured in DMEM supplemented with 25 mM glucose, 4 mM L-glutamine, 1 mM sodium pyruvate, 10% heat-inactivated FBS, and 100 units/ml penicillin–streptomycin. Cells were maintained at 37°C in a humidified 5% CO₂ incubator.

### Pharmacological reagents and calcium manipulations

To increase the intracellular calcium concentration of astrocytes, neurons, and HEK293T cells, different agents were applied: for astrocytes, TFLLR-NH2 (30 µM, Tocris); for astrocyte–neuron co-culture, KCl-intensified HEPES buffer (60 mM); and for naïve HEK293T cells, ATP-treated HEPES buffer (400 µM). For hNCX1A-expressing HEK293T cells, low-concentration sodium HEPES buffer (25 mM NaCl, 125 mM NMDG-Cl, 3 mM KCl, 2 mM CaCl₂, 2 mM MgCl₂, and 5.5 mM glucose; pH 7.3, 330 mOsmol/kg) was exchanged from normal sodium HEPES buffer (150 mM) to increase the intracellular response. To confirm the dominant role of NCX in Ca^2+^ reduction, three pharmacological agents were used: CB-DMB (1–10 µM, C5374, Sigma-Aldrich), a potent selective forward-mode NCX blocker; KB-R7943 (30 µM, 1244, Tocris), a dual-mode NCX blocker at high concentration^31,44^; and SN-6 (30 µM, 2184, Tocris), a selective reverse-mode NCX blocker. All agents were loaded for 15 min prior to delivering ultrasound stimulation.

### TAT-NCXCavBP and NCXCavBP-TAT peptides

The caveolin-binding motif in NCX1 was predicted as ΦXXXXΦXXΦ (Φ, aromatic residues; FVTAAWSIF)^64^. To generate peptides for competitive inhibition of caveolin binding, the motif was fused with the transactivating transcriptional activator (TAT) cell-penetrating peptide (YGRKKRRQRRR) at either the N- or C-terminus. TAT-conjugated CBPs (TAT-NCXCavBP: YGRKKRRQRRRFVTAAWSIF; NCXCavBP-TAT: FVTAAWSIFYGRKKRRQRRR) were synthesized and loaded into astrocytes at increasing concentrations (0.01–10 µM) before TFLLR treatment. TAT-NCXCavBP was preloaded for approximately 6 h to allow cellular uptake prior to TFLLR and ultrasound stimulation.

### NCX1 shRNA vector construction and genetic knockdown

The shRNA targeting mouse *Slc8a1* (NM_011406.3; nucleotides 421–441, 5′-CCTCTCAAGAGAAAGAAATAA-3′) was designed and synthesized as complementary oligonucleotides. The oligos were synthesized with the following structures: [phos]5′-t [sense sequence of target] ttcaagaga [reverse complement sequence of target] ttttttc-3′ and [phos]5′-tcgagaaaaaa [sense sequence of target] tctcttgaa [reverse complement sequence of target] a-3′. Two oligos were annealed with annealing buffer containing 200 mM potassium acetate, 60 mM HEPES–KOH, and 4 mM Mg-acetate (pH 7.3, adjusted using KOH), then incubated at 95°C for 5 min and 70°C for 10 min. The annealed double-stranded oligo was inserted into the HpaI-XhoI restriction enzyme sites of the pSicoR lentiviral vector (11579, Addgene) and verified by Sanger sequencing. For genetic inhibition of NCX1 in primary cultured astrocytes, 10 µg of NCX1 shRNA was electroporated (Neon Transfection system kit; MPK10096, Invitrogen) into trypsinized cultured astrocytes 4 days before the experimental day and replated onto Ultrasonocoverslips coated with 0.1 mg/ml poly-D-lysine (A-003-E, MERCK).

### Acute brain slice calcium imaging

Thy1-GCaMP6f transgenic mice (Jackson Laboratory) were used for brain slice calcium imaging. For genetic ablation, recombinant AAV vectors (serotype PHP.eB) encoding either AAV-pSicoR-Scramble sh-mCherry (6.52 × 10¹² genome copies/ml) or AAV-pSicoR-NCX1 sh-mCherry (3.61 × 10¹² genome copies/ml) were diluted 1:4 in sterile saline and administered via retro-orbital injection (20 µl total volume) into mice. Three weeks after the injection, mice were euthanized, and coronal brain slices (250 µm thick), including the ACC, were prepared using a vibratome in ice-cold ACSF with the corpus callosum serving as an anatomical landmark. Slices were allowed to recover in oxygenated ACSF at 35°C for 30 min prior to imaging. Standard ACSF composition was (in mM): 124 NaCl, 5 KCl, 1.23 NaH₂PO₄, 26 NaHCO₃, 10 glucose, 1 MgCl₂, and 2 CaCl₂. To depolarize neurons and induce calcium influx, a high-potassium ACSF solution was applied, consisting of (in mM): 64 NaCl, 60 KCl, 1.23 NaH₂PO₄, 26 NaHCO₃, 10 glucose, 1 MgCl₂, and 2 CaCl₂. All solutions were continuously bubbled with a gas mixture of 95% O₂ and 5% CO₂. Calcium imaging was performed using a Zeiss LSM 880 laser scanning confocal microscope equipped with an Airyscan detector (Carl Zeiss, Germany). Brain slices were transferred to a recording chamber and continuously superfused with ACSF at a flow rate of 3 ml/min at room temperature. GCaMP6f fluorescence was excited at 488 nm, and images were collected using a 20×/1.0 NA W Plan-Apochromat water-immersion objective. Imaging parameters were set as follows: scan area = 290.5 µm × 290.5 µm, pixel size = 1.13 µm, pixel dwell time = 2.06 µs, and frame scan time = 158.25 ms. The pinhole was adjusted to 13.47 Airy units, and the photomultiplier gain for the 488 nm channel was set to 800. Image acquisition and processing were conducted using ZEN 2.3 SP1 software (Carl Zeiss, Germany).

### RNA sequencing

For RNA sequencing analysis of NCX isoform expression, dissociated primary cortical astrocytes and neurons were prepared as described above. Cultures were harvested at 7–14 DIV (astrocytes) or 10–14 DIV (neurons) by removing the culture medium, rinsing with ice-cold phosphate-buffered saline (PBS), and lysing cells directly in TRIzol Reagent (Invitrogen). Total RNA was extracted according to the manufacturer’s protocol. RNA quality and integrity were assessed using an Agilent Bioanalyzer (RNA integrity number ≥8.0). Poly(A)-enriched mRNA libraries were prepared and sequenced on an Illumina platform. Sequencing reads were aligned to the mouse reference genome (GRCm38/mm10) using STAR. Gene expression levels were quantified as fragments per kilobase of transcript per million mapped reads (FPKM) values. NCX isoform expression (*Slc8a1*, *Slc8a2*, and *Slc8a3*) was compared between astrocytes and neurons, using *Slc6a11* (GAT3) as an astrocyte marker and *Slc17a7* (VGLUT1) as a neuronal marker.

### Structural simulation and molecular docking

To examine the effects of the D849A mutation on the NCX1 protein, we launched the simulation starting from a structure of the human NCX1 (PDB ID: 8JP0; cryo-EM structure resolution, 3.50 Å)^48^. To build up the simulation system, we used coarse-grain methods. Briefly, the experimental structure was coarse-grained through the representation of about four heavy atoms as a single bead, using Martini Bilayer Maker in CHARMM-GUI^65^. Then, the protein was embedded in a symmetric membrane consisting of 1-palmitoyl-2-oleoyl-sn-glycero-3-phosphocholine (POPC), solvated with transferable intermolecular potential with 3 points (TIP3P) water^66^ and neutralized with 0.15 M NaCl. Following equilibration, five initial poses were picked out randomly from RMSD-stabilized timepoints. These coarse-grained systems were backmapped to an all-atom system, solvated with TIP3P water, and neutralized with 0.15 M NaCl, resulting in a system containing approximately 230,000 atoms. These systems were used as the replicates for mutant generation and dynamics simulation.

To simulate the D849A mutation in the NCX1 protein, each structure was first imported into UCSF ChimeraX^67^, and the target residue, aspartate at position 849 (D849), was identified based on the corresponding sequence numbering. D849 was replaced with alanine, and the optimal conformation was selected based on the default Dunbrack rotamer library and steric compatibility^68^. Following mutation, local side-chain interactions were visually inspected to ensure the absence of steric clashes, and local energy minimization was performed using the ChimeraX Minimize Structure tool with default parameters to relieve potential steric clashes. Each structure was then solvated with TIP3P water and neutralized with 0.15 M NaCl. After equilibration, five replicates of NCX1 and five replicates of NCX1 D849A structures were subjected to production runs of 100 ns. Each production run was visualized and analyzed with UCSF ChimeraX and VMD.

### Mouse preparation for in vivo measurements

For in vivo experiments, Thy1-GCaMP6f-WPRE mice (6–10 weeks) were used regardless of sex. The calcium imaging experiments reported in this study were approved by the Institutional Animal Care and Use Committee at the Institute for Basic Science in South Korea. In behavioral experiments, male C57BL/6N mice (6–10 weeks) were used. All behavioral experiments reported in this study were approved by the Institutional Animal Care and Use Committee of Sungkyunkwan University (behavior experiments using NCX1 KD mice: SKKUIIBC2026-02-11-2, vehicle or bepridil calcium imaging experiments: SKKUIACUC2026-01-71-1, and behavior experiments with TAT-NCXCavBP peptide administration: SKKUIACUC2026-01-71-1).

### Optically transparent ultrasound transducer

An optically transparent ultrasound transducer with a 5 mm diameter and 40% optical transmittance was fabricated using PMN-PT (0.3 mm, X2B, TRS Technologies). To achieve optical transmittance and conductivity, indium-tin oxide (ITO) electrodes were deposited on both sides of the PMN-PT using a DC magnetron sputtering system (Flexlab-50, A-tech System) and annealed at 300°C for 30 s. The sample was then diced into a 3.5 mm diameter circle using a dicing machine (DAD 3221, Disco). For electrical connections between the PMN-PT and the electric wire, an inner brass housing with a 3.5 mm outer diameter and 3 mm inner diameter was adhered to the backside of the ITO-coated PMN-PT using conductive epoxy (E-solder 3022, Vonroll). This assembly was co-aligned with an outer brass housing, with a 5 mm outer diameter and 4 mm inner diameter on a glass substrate using double-sided Kapton tape. Finally, optically transparent epoxy (Epotek 301, Epoxy Technology) was carefully poured into the backside of the PMN-PT and into the gap between the inner and outer housing. The fabricated sample was then cured at room temperature for 24 h. Finally, the sample was manually polished with fine polishing compounds (05928, 3M) and poromeric polishing pads (PP8PSA, MTI Corporation) until it became optically transparent. Due to the small form factor of the transducer, a 38-µm-thick coaxial cable (38 AWG micro coaxial cable, Samtec) was precisely bonded to the exposed inner and outer housing surfaces by soldering. Subsequently, Bipolar alternating current (AC) poling was applied to the sample with a voltage of 200 V_pp_ in a square wave with 20 cycles at 1 Hz (Model 615-3, TREK)^69^. The final fabrication step involved depositing a parylene layer for electrical and water passivation using a parylene coater (OBT-PC200, Obang Technology).

### In vivo calcium imaging with odor and ultrasound stimulation

Mice were anesthetized with isoflurane (0.05%/g), and a 5-mm diameter skull cranial window was dissected at the region of an anterior–posterior (AP) +1 mm reference of bregma^52^. Following the dissection, the transparent transducer was placed over the exposed brain, and dental cement was applied along the boundary of the transducer and the remaining skull before curing it with ultraviolet light. During the surgical procedure, the body temperature of mice was maintained at 37°C using a heating pad. Following surgery, the mice were allowed to recover in a recovery room for 30 min to 1 h before starting the experiments. For wide-field calcium imaging in genetically modified mice, a lower concentration of isoflurane (0.015%/g) was utilized to avoid interference from inhalation anesthesia^70^. Calcium responses were measured using LEDs at two wavelengths with a 10-Hz frame rate on each wavelength; 470 nm light excited the GCaMP6, whereas 530 nm light was used to eliminate artifacts caused by hemodynamic changes in calcium measurements^71^. A wide-field microscope (MVX10, Olympus) was employed for calcium imaging, and the imaging trigger signals from the microscope were delivered through an Arduino board (Mega 2560, Arduino) to calculate the timing of odor and ultrasound stimulation. To deliver odor stimulation, a custom-built vaporized urine delivery system was constructed to administer mountain lion urine through a T-connection, concurrently with isoflurane to the nose mask. Ultrasound stimuli were applied 15 s after initial odor stimulation. Each experiment was performed five times on a single mouse in both vehicle and bepridil-treated groups. After completing the vehicle group experiments, the NCX inhibitor bepridil (20 mg/kg, Tocris) was administered to the mice, and subsequent experiments were conducted 1–2 h after drug administration. The vehicle solution consisted of 10% DMSO and 90% PBS. To minimize compound effects due to the presence of NCX in both the brain and heart, the drug was injected via the cisterna magna, ensuring localized delivery to the mouse brain.

### Behavioral experiments and wearable ultrasound transducer

For the predator urine exposure experiment, we monitored the freezing response of mice evoked by mountain lion urine (100% Mountain lion urine, PredatorPee). Mice were randomly assigned to independent groups for pharmacological or genetic manipulations. For NCX1 knockdown experiments, recombinant AAV-PHP.eB vectors carrying pSicoR-scramble shRNA-mCherry or pSicoR-NCX1 shRNA-mCherry were delivered into the right retro-orbital sinus under 2% isoflurane anesthesia using a 32-gauge insulin syringe. Viral stocks were diluted by mixing 10 µl of virus with 30 µl of sterile saline, and 20 µl of the diluted viral suspension was injected per mouse. Behavioral experiments were performed 2 weeks after viral injection. For the disruption of the CAV3–NCX1 interaction, either TAT-NCXCavBP (500 µM, 1 µl) or PBS (1 µl) was administered into the cisterna magna via a syringe pump at a rate of 0.1 µl/min. Mouse trajectory was recorded using a high frame rate camera (Galaxy Z Flip 3, Samsung) mounted on an XYZ stage. Prior to the experiment, a wearable ultrasonic transducer (<1 g, 5 mm diameter) with a beam collimator was attached to the exposed intact skull after removing the scalp carefully with dental cement.

The beam collimator was designed to target the ACC. Following the implantation, the mouse was moved into a cage coated with black-colored tape for recovery. Total monitoring time was 35 min, including DW exposure, urine exposure, ultrasound stimulation during urine exposure, urine-only exposure after stimulation, and 15 min of recovery in DW exposure. Kimwipes (41112, Kimtech) were separately soaked with 3 ml of DW or urine and contained in a disposable plastic dish that was replaced simultaneously with a Kimwipe to remove crosstalk effects in mouse freezing response. To further minimize such effects, all urine exposure experiments were conducted in a ventilation-enhanced room, using a separate cage for each experiment. In total, 10 shots of ultrasound stimulation were delivered with an interval of 1 min with total stimulation of 10 min. Percentage of freezing was quantified as the time during which the movement of the mouse head was less than 2 mm, relative to the total recording time. Freezing percentage was calculated as a ratio of the freezing time relative to the recording time (5 min). A wearable ultrasonic transducer was designed to be compact and lightweight, with a diameter of 5 mm and a weight of less than 1 g. A 0.3-mm thick PMN-PT (X2B, TRS Technologies) was utilized as a piezoelectric layer, and a convex PDMS lens was attached to focus the ultrasound beam at a specific depth. For the backing layer, epoxy resin (Epotek 301) was mixed with glass bubbles (K15, 3M) to minimize the transducer size as previously reported^72^. Its beam profile was measured using a needle hydrophone (NH1000, Precision Acoustics) with a step size of 50 µm in both axial and lateral directions.

### Immunoprecipitation

Intracisternal injection was performed in male C57BL/6J mice (8–10 weeks old). PBS or TAT-NCX peptide in PBS (500 µM, 1 µl) was injected into the cisterna magna at 0.1 µl/min using a Pump 11 Pico Plus Elite programmable syringe pump (Harvard Apparatus). Four hours later, the ACC was dissected and homogenized in Pierce IP Lysis Buffer. For immunoprecipitation, 2,000 µg of protein in 1 ml lysate was incubated with Protein A/G agarose (Pierce) and anti-CAV1 antibody (Invitrogen, cat. no. 03-6000, mouse monoclonal). Immunoprecipitates were analyzed by western blotting using anti-NCX1 antibody (Cell Signaling Technology, cat. no. 79350, clone D3F3H, rabbit monoclonal). Input samples contained 15 µg protein in 20 µl.

### SNI model with fiber photometry

The SNI model was established as previously described^73^. Male Thy1-GCaMP6f mice were anesthetized with 2% isoflurane and placed in the prone position. After shaving and disinfecting the right hind limb, the right sciatic nerve and its three terminal branches were exposed at the thigh level. The sural nerve was left intact, whereas the common peroneal and tibial nerves were ligated with 6-0 black silk suture and transected distal to the ligation. The wound was closed with sutures, and mice were allowed to recover before being returned to their home cage. Development of mechanical allodynia was confirmed by von-Frey testing of the ipsilateral hind paw. Behavioral, imaging, and photometry experiments were performed 7 d after SNI surgery. Age-matched non-operated mice were used for comparison. Male Thy1-GCaMP6f mice were anesthetized with 2% isoflurane and fixed in a stereotaxic frame (RWD, #68091). A cranial window (∼1 cm diameter) was prepared over the ACC region, with the dura left intact. A transparent ultrasound transducer was positioned over the window and secured with Kwik-Sil, and the optical cannula was coupled to the transducer with DW to eliminate air gaps. During recording, isoflurane was reduced to 0.25%, and body temperature was maintained at 38°C. Excitation light at 488 and 405 nm was delivered through a 400-µm optical fiber (0.4 NA, ZF1.25, Doric), and emitted fluorescence was collected through the same fiber, filtered, and detected using a fluorescence mini cube. Signals were acquired at 330 Hz. Mechanical stimulation was applied to the right hind paw using blunt forceps, and pressure-only and pressure-plus-ultrasound trials were delivered in randomized order. Ultrasound was generated using a 1.5-kHz square-wave gating signal (3.3 V_pp_, 7,500 cycles) and a 6-MHz sine-wave carrier signal (200 mV_pp_, 2,010 cycles). Signals were analyzed using custom-written scripts to calculate ΔF/F, and data from four mice were included in the statistical analysis.

### SNI model with von-Frey test and ultrasound stimulation

A total of 10 WT mice were divided into two groups: an ultrasound-treated group (n = 6; female/male = 3/3) and an SNI control group (n = 4; female/male = 2/2). The surgical procedures for the SNI model were identical to those described for the fiber photometry experiments. In TAT-NCXCavBP application experiments, TAT-NCXCavBP peptide (YGRKKRRQRRRFVTAAWSIF), consisting of the TAT cell-penetrating sequence fused to the NCX1 caveolin-binding motif, was administered into the cisterna magna. Mice were anesthetized with 2% isoflurane, and an incision was made in the skin over the back of the neck. The peptide solution was injected into the cisterna magna using a 32-gauge insulin syringe needle. Each mouse received 1.3 µg of peptide in 1.3 µl of solution at a concentration of 1 mg/ml. The injection rate was 0.1 µg/min (0.1 µl/min), giving a total injection time of 13 min. After injection, the incision site was disinfected and sutured. Injections were performed on postoperative days 10, 12, 14, 16, and 18, with a day between procedures to allow recovery. Six hours after the start of peptide administration, mice in the ultrasound-treated group were anesthetized and head-fixed. Isoflurane was maintained at 0.2% during stimulation, and 6-MHz ultrasound was applied to the ACC for 5 s. Ultrasound stimulation was performed on each peptide administration day. The peptide-only group received peptide injections on the same schedule without ultrasound stimulation.

Mechanical sensitivity was assessed using the von-Frey test. Mice were placed individually in transparent acrylic chambers on a metal platform and allowed to acclimate for at least 1 h before testing. Paw withdrawal thresholds of the ipsilateral hind paw were measured using calibrated von-Frey filaments. Filaments were applied to the plantar surface of the ipsilateral hind paw, and a brisk paw withdrawal, lifting, or flinching response was considered a positive response. The 50% paw withdrawal threshold was calculated using the up-down method as previously described^74^. The von-Frey tests were performed before injury and on postoperative days 2, 4, 8, 10–14, 16, 18, and 21, and 6-MHz ultrasound stimulation was applied for 5 s from postoperative day 10 to 14. The SNI control group underwent the same surgical and behavioral testing procedures without ultrasound stimulation. In the TAT-NCXCavBP experiment, mechanical withdrawal thresholds were measured on postoperative days 2, 4, and 8 to assess the development of mechanical allodynia. Further testing was performed on treatment days 10, 12, 14, 16, 18, and 20, with a final follow-up measurement on day 23.

### ECoG recording in the murine model of KA-induced seizures

A total of 13 male mice were used and divided into two groups: ultrasound-treated group (n = 9) and KA control group (n = 4). Mice were anesthetized with 2% isoflurane and placed in a stereotaxic frame. After exposure of the skull, burr holes were made at AP −2.1 mm, ML −1.6 mm relative to bregma using a 1-mm drill bit. Stainless steel screw electrodes were then implanted for ECoG recording. KA was dissolved in 0.9% sterile NaCl, pH 7.4, at a concentration of 20 mM. For intrahippocampal KA injection, a Nanoject glass capillary (3.5-inch length; Drummond Scientific, 3-000-203-G/X) was connected to a Picospritzer and lowered into the right dorsal hippocampal CA1 region at AP −2.1 mm, ML +1.6 mm, and DV −1.7 mm relative to bregma. A total volume of 100 nl KA solution was delivered slowly over 10 min by pressure injection. After injection, the capillary was left in place for an additional 10 min to minimize backflow and was then slowly withdrawn. Ultrasound (6 MHz) was applied for 5 s per trial over 10 trials at 1 min intervals, giving a total stimulation period of 10 min. For ECoG spike counting, a threshold of 2.5 standard deviations of the signal was applied. Spike frequencies were determined per minute across matched time intervals for both US-treated and KA control groups. The change in spike frequency was calculated by subtracting the baseline-normalized frequency of the KA control group from the US-treated group.

### Immunohistochemistry

Mice were transcardially perfused with PBS followed by 4% paraformaldehyde (PFA), and extracted brains were post-fixed in 4% PFA overnight. The tissue was then sectioned at 30 µm into cold PBS using a vibratome (Leica VT1000S) and stored at 4°C. For c-Fos and NeuN labeling, sections were blocked with 5% normal goat serum in PBS/0.3% Triton X-100 (PBST) at room temperature for 1 h and incubated overnight with rabbit anti-c-Fos (1:1,000, SYSY-226008) and mouse anti-NeuN (1:1,000, Millipore MAB377) antibodies. After PBS washes, sections were incubated with Alexa Fluor 568-conjugated anti-rabbit (Abcam ab175471, 1:800) and Alexa Fluor 488-conjugated anti-mouse secondary antibodies (Abcam ab150113, 1:800). Sections were washed with PBST four times before they were mounted onto glass slides and coverslipped with DAPI-containing mounting medium VECTASHIELD (Vector Laboratories). Imaging quantifications of c-Fos and NeuN were conducted at 20× on AxioScan Z1. For quantitative analysis, c-Fos^+^/NeuN⁺ cells were counted as an average of three sections per animal.

### Immunoblotting

Cultured astrocytes were harvested and homogenized with a pestle in Tissue Lysis/Extraction Reagent (Sigma-Aldrich). For brain tissue samples, ACC or whole-cortex regions were microdissected from liquid nitrogen-frozen coronal brain sections and homogenized in PRO-PREP Protein Extraction Solution for Cell/Tissue (iNtRON Biotechnology). Cleared lysates were quantified using a BCA assay, and 15 µg of total protein per lane was resolved on hand-cast 8% Tris–glycine gels and transferred to PVDF membranes. Astrocyte membranes were blocked in 5% nonfat milk, whereas brain tissue membranes were blocked in 5% BSA and incubated overnight at 4°C with primary antibodies diluted in 1% BSA. Primary antibodies were rabbit anti-NCX1 (1:1,000; Abcam, ab151608), rabbit anti-NCX2 (1:1,000; Alomone Labs, ANX-012), rabbit anti-NCX3 (1:10,000; Alomone Labs, ANX-013), and rabbit anti-GAPDH (1:5,000; Cell Signaling Technology, 2118S). Membranes were incubated with HRP-conjugated anti-rabbit secondary antibody, and chemiluminescent signals were imaged using Image Lab and quantified with ImageJ. Protein levels were normalized to GAPDH levels.

### Statistics

Data are presented as the mean ± standard error of the mean. The normality of the data distribution was assessed using the Shapiro–Wilk test. For comparison between two groups, a paired or unpaired t-test was employed when the data followed a normal distribution; otherwise, an appropriate nonparametric test was used. For comparisons among multiple groups, one-way or two-way ANOVA was performed, followed by post hoc multiple comparison tests tailored to the normality of variance. Significance was defined as *p* < 0.05 (\**p* < 0.05, \*\**p* < 0.01, \*\*\**p* < 0.001, ns: not significant). Exact *P*-values and sample sizes are reported in the figure legends. All analyses were performed using GraphPad Prism.

## Data availability

Data and code are available from the corresponding authors upon reasonable request.

## Author contributions

J.P., J.M.P., and K.L. developed the conception and wrote the original draft. K.L. fabricated all of the ultrasonic transducers, performed all in vitro experiments, including astrocyte, astrocyte– neuron co-culture, pharmacology, genetic knockdown, genetic overexpression, TAT peptide experiments, and in vivo experiments, including odor exposure calcium imaging, pharmacology, and behavior experiments. J.M.L. performed in vitro experiments, including astrocyte, astrocyte– neuron co-culture, pharmacology, genetic knockdown, genetic overexpression, designed TAT peptide, knockdown, overexpression models in vitro and in vivo. H.K. and S.L. performed TAT peptide in vitro experiments, western blot analysis, co-immunoprecipitation analysis, MD simulation, and SNI and ECoG in vivo experiments. M.K. fabricated all of the ultrasonic transducers, performed genetic knockdown, genetic overexpression, TAT peptide in vitro experiments, ECoG in vivo experiments, and in vivo behavior experiments. S.S. performed acute brain slice calcium imaging experiments. W.Y. performed MD simulation. T.T.P. performed immunohistochemistry and western blot analysis. C.J.L., J.P., and J.M.P. supervised all of the experiments. K.L., H.K., C.J.L., J.P., and J.M.P. reviewed and edited the manuscript.

## References

1. Raza, M., et al. Evidence that injury-induced changes in hippocampal neuronal calcium dynamics during epileptogenesis cause acquired epilepsy. Proc. Natl Acad. Sci. USA 101, 17522–17527 (2004).

2. Delorenzo, R. J., Sun, D. A. & Deshpande, L. S. Cellular mechanisms underlying acquired epilepsy: the calcium hypothesis of the induction and maintainance of epilepsy. Pharmacol. Ther. 105, 229–266 (2005).

3. Deuschl, G. et al. A randomized trial of deep-brain stimulation for Parkinson’s disease. N. Engl. J. Med. 355, 896–908 (2006).

4. Boon, P. et al. Deep brain stimulation in patients with refractory temporal lobe epilepsy. Epilepsia 48, 1551–1560 (2007).

5. Vitek, J. L. Mechanisms of deep brain stimulation: excitation or inhibition. Mov. Disord. 17, S69–S72 (2002).

6. Nguyen, J. P., Nizard, J., Keravel, Y. & Lefaucheur, J. P. Invasive brain stimulation for the treatment of neuropathic pain. Nat. Rev. Neurol. 7, 699–709 (2011).

7. Lozano, A. M. & Lipsman, N. Probing and regulating dysfunctional circuits using deep brain stimulation. Neuron 77, 406–424 (2013).

8. Silvanto, J., Muggleton, N. & Walsh, V. State-dependency in brain stimulation studies of perception and cognition. Trends Cogn. Sci. 12, 447–454 (2008).

9. Bradley, C., Nydam, A. S., Dux, P. E. & Mattingley, J. B. State-dependent effects of neural stimulation on brain function and cognition. Nat. Rev. Neurosci. 23, 459–475 (2022).

10. Rabut, C. et al. Ultrasound technologies for imaging and modulating neural activity. Neuron 108, 93–110 (2020).

11. Tufail, Y., Yoshihiro, A., Pati, S., Li, M. M. & Tyler, W. J. Ultrasonic neuromodulation by brain stimulation with transcranial ultrasound. Nat. Protoc. 6, 1453–1470 (2011).

12. Choi, T. et al. Bidirectional neuronal control of epileptiform activity by repetitive transcranial focused ultrasound stimulations. Adv. Sci. 11, e2302404 (2024).

13. Murphy, K. R. et al. Optimized ultrasound neuromodulation for non-invasive control of behavior and physiology. Neuron 112, 3252–3266.e5 (2024).

14. Oh, S. J. et al. Ultrasonic neuromodulation via astrocytic TRPA1. Curr. Biol. 29, 3386–3401.e8 (2019).

15. Lee, K. et al. Ultrasonocoverslip: in-vitro platform for high-throughput assay of cell type-specific neuromodulation with ultra-low-intensity ultrasound stimulation. Brain Stimul. 16, 1533–1548 (2023).

16. Yoo, S., Mittelstein, D. R., Hurt, R. C., Lacroix, J. & Shapiro, M. G. Focused ultrasound excites cortical neurons via mechanosensitive calcium accumulation and ion channel amplification. Nat. Commun. 13, 493 (2022).

17. Qiu, Z. et al. The mechanosensitive ion channel Piezo1 significantly mediates in vitro ultrasonic stimulation of neurons. iScience 21, 448–457 (2019).

18. Matsushita, Y., et al. TRPC6 is a mechanosensitive channel essential for ultrasound neuromodulation in the mammalian brain. Proc. Natl Acad. Sci. USA 121, e2404877121 (2024).

19. Duque, M. et al. Sonogenetic control of mammalian cells using exogenous Transient Receptor Potential A1 channels. Nat. Commun. 13, 600 (2022).

20. Sorum, B., et al. Ultrasound activates mechanosensitive TRAAK K+ channels through the lipid membrane. Proc. Natl Acad. Sci. USA 118, e2006980118 (2021).

21. Reeves, J. P., Abdellatif, M. & Condrescu, M. The sodium-calcium exchanger is a mechanosensitive transporter. J. Physiol. 586, 1549–1563 (2008).

22. Dong, H., Tang, B., Jiang, Y. & Mittal, R. K. Na+/Ca2+ exchanger 1 is a key mechanosensitive molecule of the esophageal myenteric neurons. Acta Physiol. 225, e13223 (2019).

23. Bossuyt, J., Taylor, B. E., James-Kracke, M. & Hale, C. C. The cardiac sodium-calcium exchanger associates with caveolin-3. Ann. NY Acad. Sci. 976, 197–204 (2002).

24. Lang, D. et al. Caveolar compartmentalization of pacemaker signaling ensures stable sinoatrial rhythmicity which is disrupted in heart failure. JACC Clin. Electrophysiol. 12, 957–976 (2026).

25. Sedding, D. G. et al. Caveolin-1 facilitates mechanosensitive protein kinase B (Akt) signaling in vitro and in vivo. Circ. Res. 96, 635–642 (2005).

26. Yang, H. et al. Mechanosensitive caveolin-1 activation-induced PI3K/Akt/mTOR signaling pathway promotes breast cancer motility, invadopodia formation and metastasis in vivo. Oncotarget 7, 16227–16247 (2016).

27. Dallapiazza, R. F. et al. Noninvasive neuromodulation and thalamic mapping with low-intensity focused ultrasound. J. Neurosurg. 128, 875–884 (2018).

28. Wang, F. et al. Astrocytes modulate neural network activity by Ca2+-dependent uptake of extracellular K+. Sci. Signal. 5, ra26 (2012).

29. Shehata, M., Matsumura, H., Okubo-Suzuki, R., Ohkawa, N. & Inokuchi, K. Neuronal stimulation induces autophagy in hippocampal neurons that is involved in AMPA receptor degradation after chemical long-term depression. J. Neurosci. 32, 10413–10422 (2012).

30. Diallinas, G. Understanding transporter specificity and the discrete appearance of channel-like gating domains in transporters. Front. Pharmacol. 5, 207 (2014).

31. Dubyak, G. R. Ion homeostasis, channels, and transporters: an update on cellular mechanisms. Adv. Physiol. Educ. 28, 143–154 (2004).

32. Hilgemann, D. W. Cytoplasmic ATP-dependent regulation of ion transporters and channels: mechanisms and messengers. Annu. Rev. Physiol. 59, 193–220 (1997).

33. Al Awabdh, S. et al. Neuronal activity mediated regulation of glutamate transporter GLT-1 surface diffusion in rat astrocytes in dissociated and slice cultures. Glia 64, 1252–1264 (2016).

34. Liu, B., Peel, S. E., Fox, J. & Hall, I. P. Reverse mode Na^+^/Ca^2+^ exchange mediated by STIM1 contributes to Ca^2+^ influx in airway smooth muscle following agonist stimulation. Respir. Res. 11, 168 (2010).

35. Clapham, D. E. Calcium signaling. Cell 131, 1047–1058 (2007).

36. Blaustein, M. P. & Lederer, W. J. Sodium/calcium exchange: its physiological implications. Physiol. Rev. 79, 763–854 (1999).

37. Deitmer, J. W., Verkhratsky, A. J. & Lohr, C. Calcium signalling in glial cells. Cell Calcium 24, 405–416 (1998).

38. Rose, C. R., Ziemens, D. & Verkhratsky, A. On the special role of NCX in astrocytes: Translating Na^+^-transients into intracellular Ca^2+^ signals. Cell Calcium 86, 102154 (2020).

39. Reyes, R. C., Verkhratsky, A. & Parpura, V. Plasmalemmal Na^+^/Ca^2+^ exchanger modulates Ca^2+^-dependent exocytotic release of glutamate from rat cortical astrocytes. ASN Neuro 4, e00075 (2012).

40. Liu, Z., Cheng, Q., Ma, X. & Song, M. Suppressing effect of Na^+^/Ca^2+^ exchanger (NCX) inhibitors on the growth of melanoma cells. Int. J. Mol. Sci. 23, 901 (2022).

41. Hu, H. J. et al. Blockade of the forward Na^+^/Ca^2+^ exchanger suppresses the growth of glioblastoma cells through Ca^2+^-mediated cell death. Br. J. Pharmacol. 176, 2691–2707 (2019).

42. Iwamoto, T. et al. The exchanger inhibitory peptide region-dependent inhibition of Na^+^/Ca^2+^ exchange by SN-6 [2-[4-(4-nitrobenzyloxy)benzyl]thiazolidine-4-carboxylic acid ethyl ester], a novel benzyloxyphenyl derivative. Mol. Pharmacol. 66, 45–55 (2004).

43. Liang, G. H., Kim, J. A., Seol, G. H., Choi, S. & Suh, S. H. The Na^+^/Ca^2+^ exchanger inhibitor KB-R7943 activates large-conductance Ca^2+^-activated K^+^ channels in endothelial and vascular smooth muscle cells. Eur. J. Pharmacol. 582, 35–41 (2008).

44. Watano, T., Kimura, J., Morita, T. & Nakanishi, H. A novel antagonist, No. 7943, of the Na^+^/Ca^2+^ exchange current in guinea-pig cardiac ventricular cells. Br. J. Pharmacol. 119, 555–563 (1996).

45. Thurneysen, T., Nicoll, D. A., Philipson, K. D. & Porzig, H. Sodium/calcium exchanger subtypes NCX1, NCX2 and NCX3 show cell-specific expression in rat hippocampus cultures. Brain Res. Mol. Brain Res. 107, 145–156 (2002).

46. Canitano, A. et al. Brain distribution of the Na^+^/Ca^2+^ exchanger-encoding genes NCX1, NCX2, and NCX3 and their related proteins in the central nervous system. Ann. N. Y. Acad. Sci. 976, 394–404 (2002).

47. Yang, Y. C. et al. Regulation of sodium-calcium exchanger activity by creatine kinase under energy-compromised conditions. J. Biol. Chem. 285, 28275–28285 (2010).

48. Dong, Y. et al. Structural insight into the allosteric inhibition of human sodium-calcium exchanger NCX1 by XIP and SEA0400. EMBO J. 43, 2 (2024).

49. Whiteley, G., Collins, R. F. & Kitmitto, A. Characterization of the molecular architecture of human caveolin-3 and interaction with the skeletal muscle ryanodine receptor. J. Biol. Chem. 287, 40302–40316 (2012).

50. Couet, J., Li, S., Okamoto, T., Ikezu, T. & Lisanti, M. P. Identification of peptide and protein ligands for the caveolin-scaffolding domain. Implications for the interaction of caveolin with caveolae-associated proteins. J. Biol. Chem. 272, 6525–6533 (1997).

51. Collins, B. M., Davis, M. J., Hancock, J. F. & Parton, R. G. Structure-based reassessment of the caveolin signaling model: do caveolae regulate signaling through caveolin-protein interactions? Dev. Cell 23, 11–20 (2012).

52. Jhang, J. et al. Anterior cingulate cortex and its input to the basolateral amygdala control innate fear response. Nat. Commun. 9, 2744 (2018).

53. Wernecke, K. E. et al. Fox urine exposure induces avoidance behavior in rats and activates the amygdalar olfactory cortex. Behav. Brain Res. 279, 76–81 (2015).

54. Pan, G., Yang, J. M., Hu, X. Y. & Li, X. M. Postnatal development of the electrophysiological properties of somatostatin interneurons in the anterior cingulate cortex of mice. Sci. Rep. 6, 28137 (2016).

55. Coste, B. et al. Piezo1 and Piezo2 are essential components of distinct mechanically activated cation channels. Science 330, 55–60 (2010).

56. Wang, Y. Y., Chang, R. B., Waters, H. N., McKemy, D. D. & Liman, E. R. The nociceptor ion channel TRPA1 is potentiated and inactivated by permeating calcium ions. J. Biol. Chem. 283, 32691–32703 (2008).

57. Polat, O. K. et al. Contribution of coiled-coil assembly to Ca^2+^/calmodulin-dependent inactivation of TRPC6 channel and its impacts on FSGS-associated phenotypes. J. Am. Soc. Nephrol. 30, 1587–1603 (2019).

58. Sinha, B. et al. Cells respond to mechanical stress by rapid disassembly of caveolae. Cell 144, 402–413 (2011).

59. Pinton, G. F. et al. Attenuation, scattering, and absorption of ultrasound in the skull bone. Med. Phys. 39, 299–307 (2012).

60. Murphy, K. R. et al. A practical guide to transcranial ultrasonic stimulation from the IFCN-endorsed ITRUSST consortium. Clin. Neurophysiol. 171, 192–226 (2025).

61. Aubry, J. F. et al. ITRUSST consensus on biophysical safety for transcranial ultrasound stimulation. Brain Stimul. 18, 1896–1905 (2025).

62. Jo, Y., Kim, S., Jeong, J. & Lee, H. J. Ultrasound brain stimulation technologies for targeted therapeutics. Nat. Electron. 8, 647–662 (2025).

63. Sarvazyan, A. P., Rudenko, O. V. & Fatemi, M. Acoustic radiation force: A review of four mechanisms for biomedical applications. IEEE Trans. Ultrason. Ferroelectr. Freq. Control 68, 3261–3269 (2021).

## References

64. Okamoto, T., Schlegel, A., Scherer, P. E. & Lisanti, M. P. Caveolins, a Family of Scaffolding Proteins for Organizing “Preassembled Signaling Complexes” at the Plasma Membrane. J. Biol. Chem. 273, 5419–5422 (1998).

65. Jo, S., Kim, T., Iyer, V. G. & Im, W. CHARMM-GUI: a web-based graphical user interface for CHARMM. J. Comput. Chem. 29, 1859–1865 (2008).

66. Jorgensen, W. L., Chandrasekhar, J., Madura, J. D., Impey, R. W. & Klein, M. L. Comparison of simple potential functions for simulating liquid water. J. Chem. Phys. 79, 926–935 (1983).

67. Pettersen, E. F. et al. UCSF ChimeraX: Structure visualization for researchers, educators, and developers. Protein Sci. 30, 70–82 (2021).

68. Dunbrack, R. L. Jr., Rotamer libraries in the 21st century. Curr. Opin. Struct. Biol. 12, 431–440 (2002).

69. Qiu, C. et al. Transparent ferroelectric crystals with ultrahigh piezoelectricity. Nature 577, 350–354 (2020).

70. Yuan, Y., Wang, X., Yan, J. & Li, X. The effect of anesthetic dose on the motor response induced by low-intensity pulsed ultrasound stimulation. BMC Neurosci. 19, 78 (2018).

71. Ma, Y. et al. Wide-field optical mapping of neural activity and brain haemodynamics: considerations and novel approaches. Philos. Trans. R. Soc. Lond. B Biol. Sci. 371, 20150360 (2016).

72. Kim, H. et al. High-attenuation backing layer for miniaturized ultrasound imaging transducer. IEEE Trans. Ultrason. Ferroelectr. Freq. Control 69, 1960–1969 (2022).

73. Decosterd, I. & Woolf, C. J. Spared nerve injury: an animal model of persistent peripheral neuropathic pain. Pain 87, 149–158 (2000).

74. Chaplan, S. R., Bach, F. W., Pogrel, J. W., Chung, J. M. & Yaksh, T. L. Quantitative assessment of tactile allodynia in the rat paw. J. Neurosci. Methods 53, 55–63 (1994).

