## Supplementary Information for "Selective ultrasonic suppression of pathological brain hyperactivity via an endogenous CAV3–NCX1 axis"

**This file includes:**

Supplementary Figs. 1–11

### Supplementary Figures

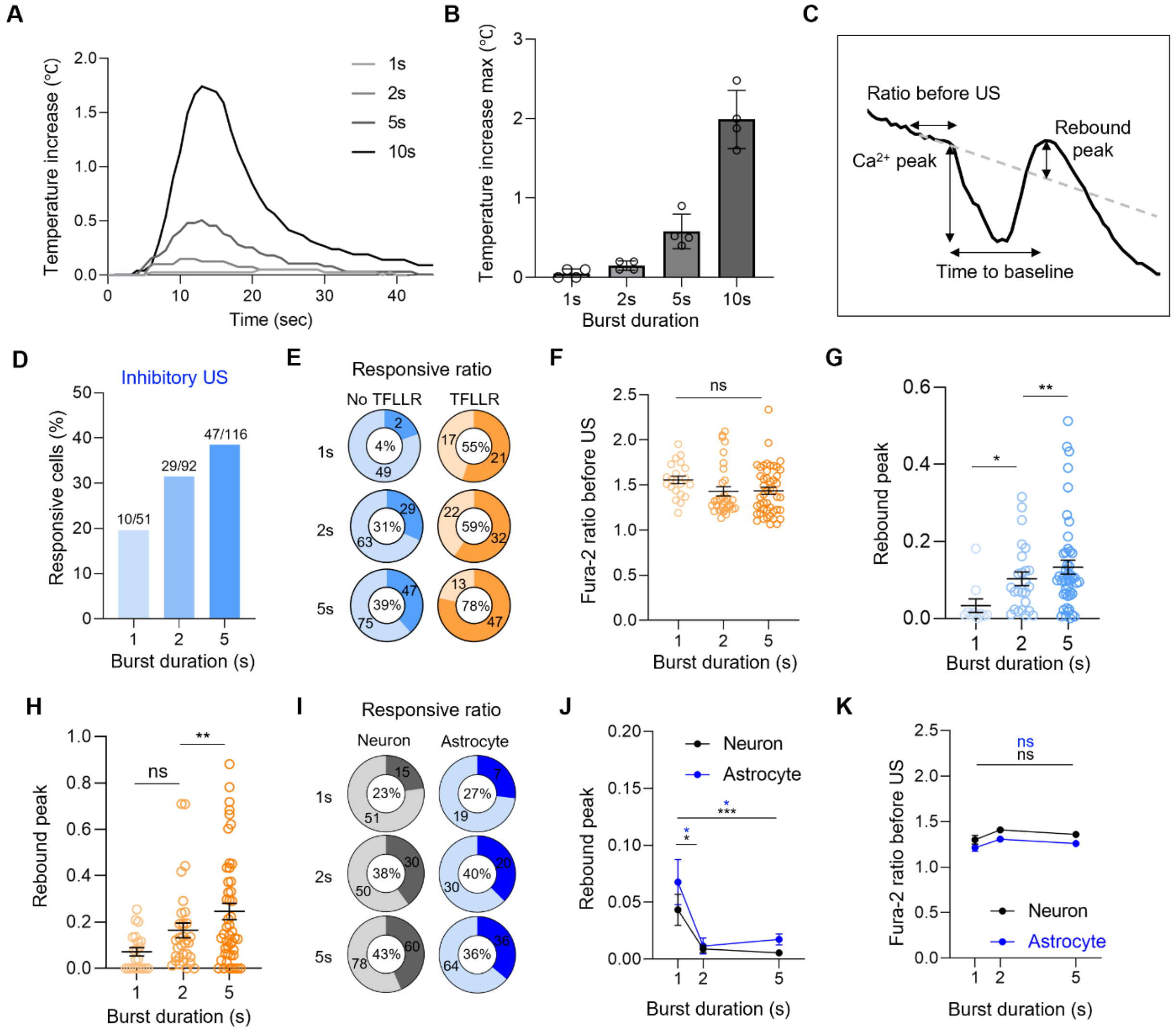


**Supplementary Fig. 1 Thermal effects and cellular response profiles for ultrasound stimulation of different burst durations. a**, Temperature increase (°C) over time on the transducer surface during ultrasound stimulation at four burst durations (1, 2, 5, and 10 s). **b**, Maximum temperature increase (°C) as a function of burst duration. **c**, Diagram of a representative $Ca^{2+}$ trace illustrating the four key quantitative parameters: $Ca^{2+}$ peak amplitude, Fura-2 ratio before ultrasound (US, pre-stimulus baseline), rebound peak, and time to baseline. **d**, Percentage of responsive cells as a function of ultrasound burst duration upon inhibitory stimulation. n = 51, 92, and 122 for 1 s, 2 s, and 5 s, respectively. **e**, Pie charts showing the proportions of ultrasound-responsive cells in No-TFLLR (left) and TFLLR-treated (right) conditions across burst durations. **f**, Fura-2 ratio before ultrasound stimulation across burst durations in TFLLR-treated cultured astrocytes. Kruskal–Wallis test with Dunn's multiple comparison test. **g**, Rebound peak amplitude in cultured astrocytes across burst durations without TFLLR application. Kruskal–Wallis test with Dunn's multiple comparison test. **h**, Rebound peak amplitudes across burst durations in TFLLR-treated cultured astrocytes. A duration-dependent increase in rebound peak is observed. Kruskal–Wallis test with Dunn's multiple comparison test. **i**, Pie charts showing the proportions of ultrasound-responsive cells among neurons (red) and astrocytes (gray) across burst durations in KCl-treated co-cultures. **j**, Rebound peak amplitude comparing neurons (red) and astrocytes (black) across burst durations in KCl-treated co-cultures. The amplitude increases as a function of burst duration. Kruskal–Wallis test with Dunn's multiple comparison test. **k**, Fura-2 ratio before ultrasound stimulation of neurons (red) and astrocytes (black) across burst durations in KCl-treated co-cultures. Kruskal–Wallis test with Dunn's multiple comparison test.

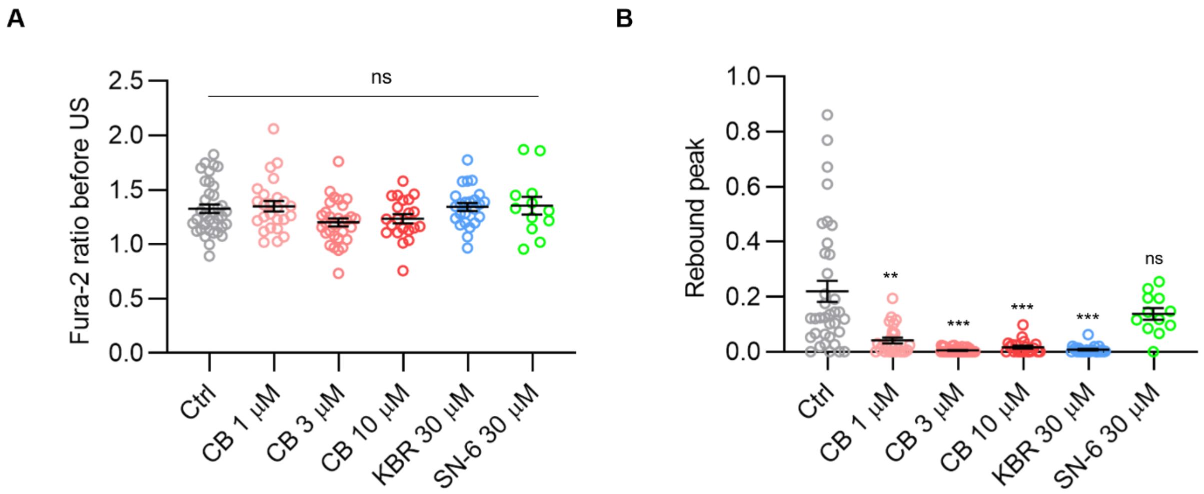


**Supplementary Fig. 2 Control experiments for NCX mode-specific drugs. a**, Pre-stimulation Fura-2 ratio before ultrasound (US) onset across pharmacological conditions, confirming comparable intracellular $Ca^{2+}$ levels at baseline across all drug treatment groups. Kruskal–Wallis test. **b**, Rebound peak amplitude following the ultrasound-induced suppression of $Ca^{2+}$ signals across pharmacological conditions. Brown–Forsythe and Welch ANOVA with Dunnett's multiple comparison test.

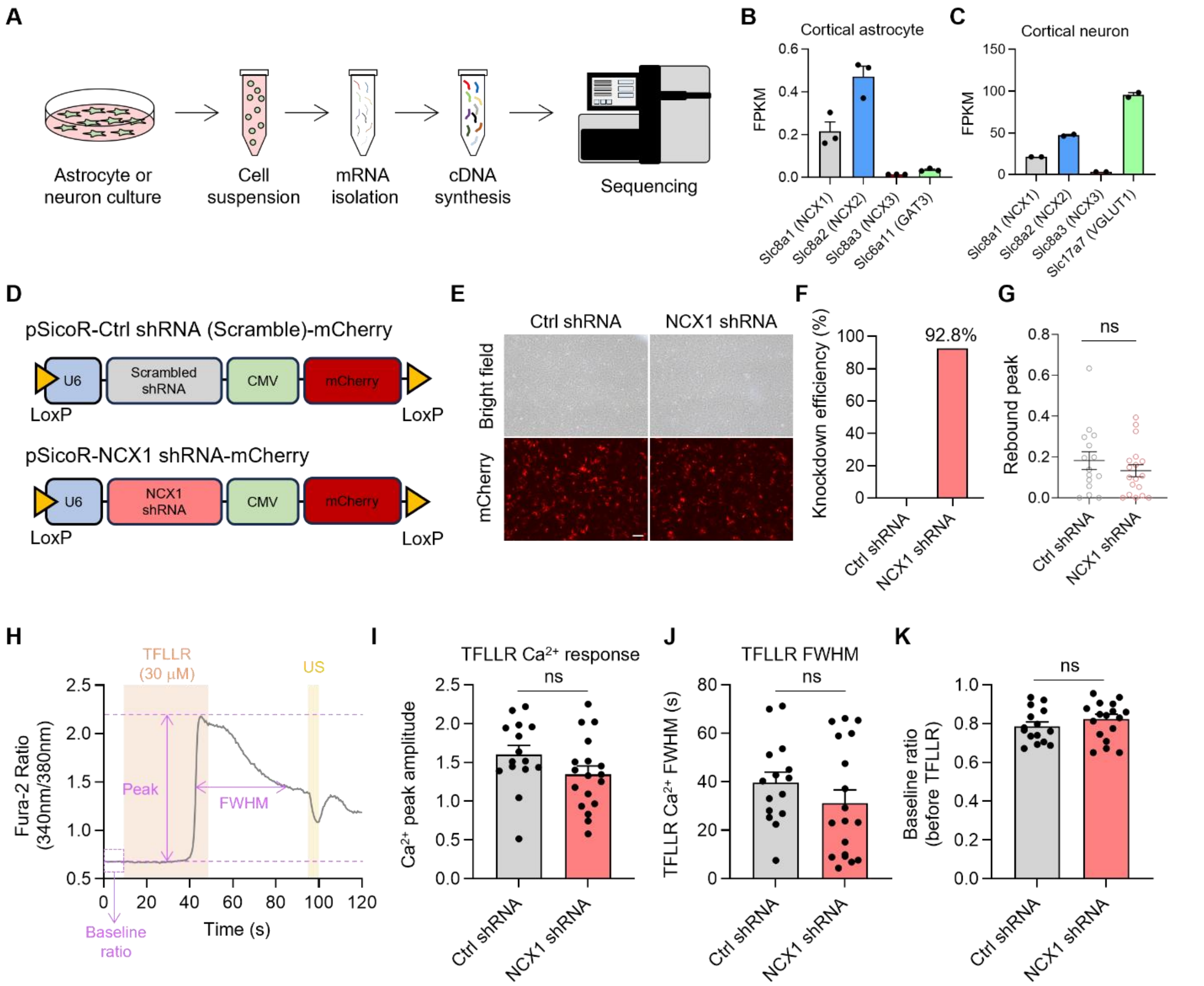


**Supplementary Fig. 3 NCX1 shRNA construct, knockdown efficiency, and baseline TFLLR calcium response.** **a**, Workflow for RNA sequencing of cultured cortical astrocytes and neurons. Dissociated cultures were prepared as single-cell suspensions, followed by mRNA isolation, cDNA library synthesis, and high-throughput sequencing. **b**, FPKM values for *Slc8a1* (NCX1), *Slc8a2* (NCX2), *Slc8a3* (NCX3), and *Slc6a11* (GAT3, astrocyte marker) in cultured cortical astrocytes. **c**, FPKM values for *Slc8a1* (NCX1), *Slc8a2* (NCX2), *Slc8a3* (NCX3), and *Slc17a7* (VGLUT1, neuronal marker) in cultured cortical neurons. **d**, Schematic of shRNA construct design. pSicoR-Ctrl shRNA (Scramble)-mCherry (top): U6-driven scrambled shRNA with CMV-driven mCherry reporter. pSicoR-NCX1 shRNA-mCherry (bottom): identical architecture with NCX1-targeting shRNA sequence. **e**, Representative brightfield and mCherry fluorescence images of astrocytes transfected with Ctrl shRNA or NCX1 shRNA, confirming efficient transfection. **f**, Knockdown efficiency of NCX1 protein following NCX1 shRNA transfection in cultured astrocytes, assessed by western blot and normalized to the expression under Ctrl shRNA conditions. NCX1 shRNA achieves 92.8% knockdown efficiency. **g**, Rebound peak amplitude following the ultrasound-induced suppression of $Ca^{2+}$ levels in astrocytes transfected with Ctrl shRNA or NCX1 shRNA. Mann–Whitney unpaired test. **h**, Representative Fura-2 ratio time-course trace showing TFLLR application and subsequent ultrasound stimulation. **i**, TFLLR $Ca^{2+}$ peak amplitude comparing astrocytes transfected with Ctrl shRNA or NCX1 shRNA. Unpaired t-test. **j**, TFLLR $Ca^{2+}$ transient full with at half maximum (FWHM) comparing astrocytes transfected with Ctrl shRNA or NCX1 shRNA. Mann–Whitney unpaired test. **k**, Baseline Fura-2 ratio before TFLLR application in astrocytes transfected with Ctrl shRNA or NCX1 shRNA. Mann–Whitney unpaired test.

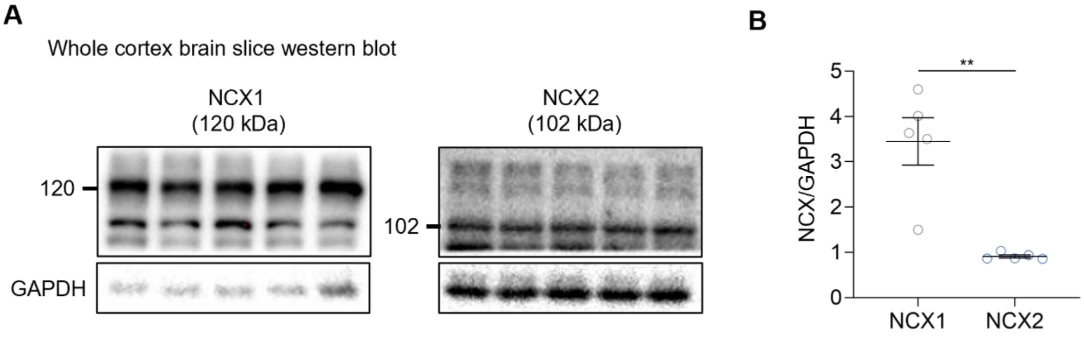


**Supplementary Fig. 4 NCX isoform expression in the murine cortex and overview of isoform selectivity. a**, Western blot comparing NCX1 (120 kDa) and NCX2 (102 kDa) protein expression in whole-cortex brain slice lysates from adult mice. GAPDH (36 kDa) serves as a loading control. **b**, Quantification of NCX1 and NCX2 protein levels normalized to GAPDH levels in whole-cortex lysates. n = 5 per group. Unpaired t-test.

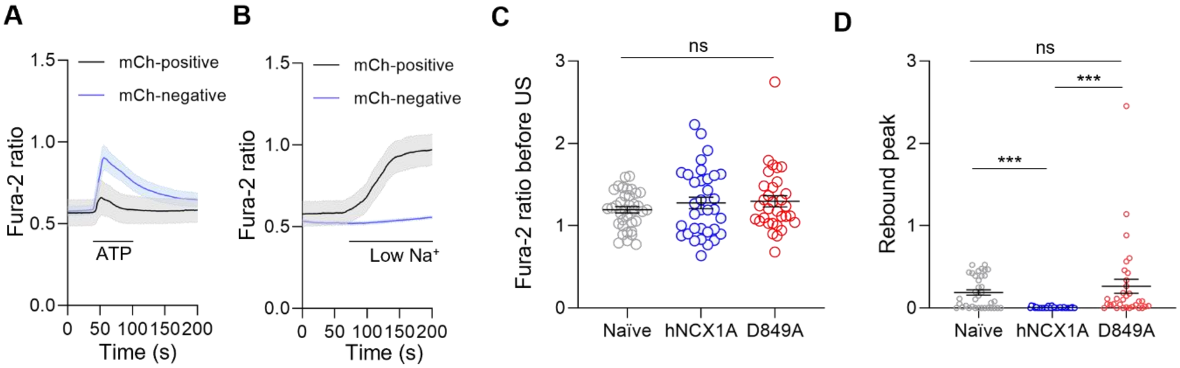


**Supplementary Fig. 5 Validation of hNCX1A overexpression and comparison of Fura-2 ratios before ultrasound stimulation and during the rebound peak in hNCX1A- and D849A mutant-expressing cells. a**, Fura-2 ratio time course in mCherry (mCh)-positive (black, hNCX1A-expressing) and mCherry-negative (blue, naïve) HEK293T cells during ATP application. **b**, Fura-2 ratio time course in mCherry-positive (black, hNCX1A-expressing) and mCherry-negative (blue, naïve) HEK293T cells during the application of low-$Na^+$ solution. **c**, Pre-stimulation Fura-2 ratio before ultrasound among naïve, mutant-expressing, and D849A mutant-expressing HEK293T cells, confirming comparable intracellular $Ca^{2+}$ levels at baseline. ns, Kruskal–Wallis test with Dunn's multiple comparison. **d**, Rebound peak amplitude following the ultrasound-induced suppression of $Ca^{2+}$ levels among naïve, mutant-expressing, and D849A mutant-expressing HEK293T cells. Kruskal–Wallis test with Dunn's multiple comparison.

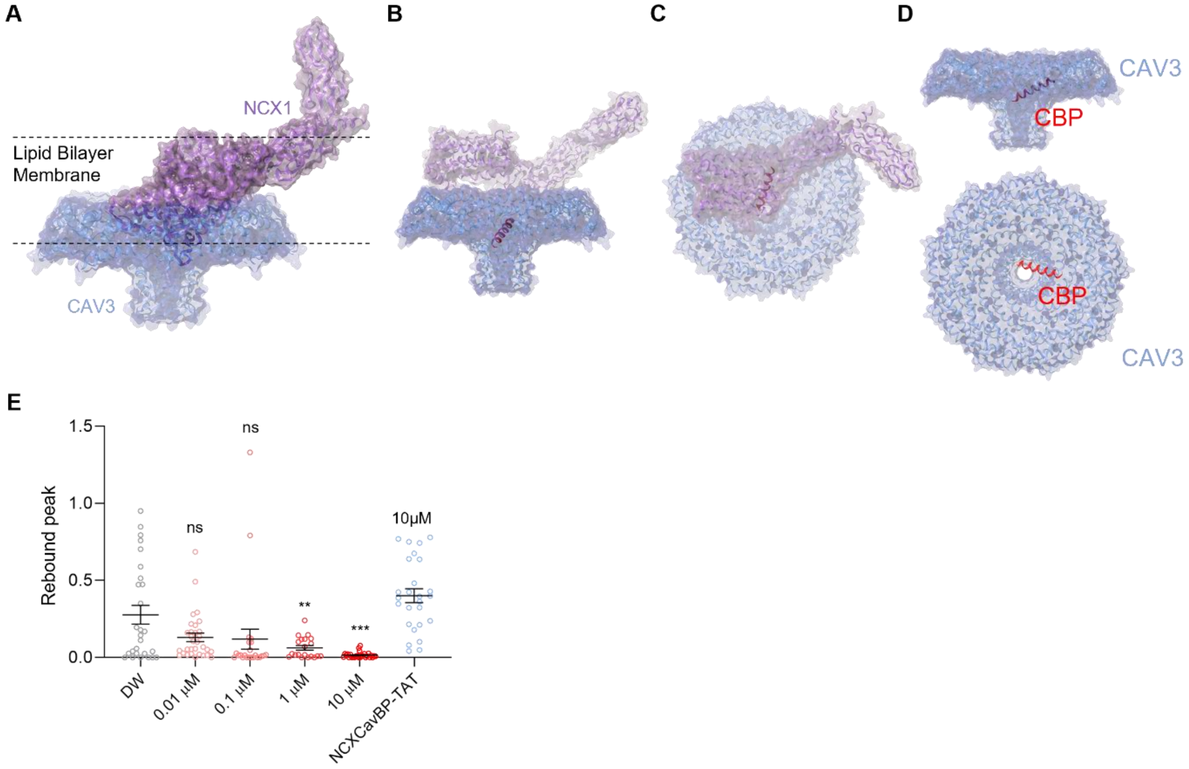


**Supplementary Fig. 6 Multi-perspective views of the NCX1–CAV3 complex highlighting the CBP binding interface. a**, Side view of the structural docking model of NCX1 (purple, surface representation) positioned atop the CAV3 nonamer (blue, surface representation) at the lipid bilayer membrane. Dashed lines indicate the upper and lower boundaries of the lipid bilayer. Structural models generated using AlphaFold2. Docking performed using Schrödinger PIPER software. **b**, Alternative, rotated side view of the NCX1–CAV3 complex docking model. **c**, Top-down (extracellular) view of the CAV3–NCX1 complex, showing the spatial relationship between NCX1 and the CAV3 nonamer ring scaffold. **d**, Two structural views (side view, left; top-down view, right) of the caveolin-binding peptide (CBP, red helical ribbon) bound to the CAV3 nonamer (blue surface). The CBP occupies the NCX1-binding surface at the CAV3 scaffolding domain. **e**, Rebound peak amplitude following ultrasound stimulation across TAT-NCXCavBP conditions in TFLLR-treated astrocytes. Vehicle (DW), TAT-NCXCavBP 0.01 μM, TAT-NCXCavBP 0.1 μM, TAT-NCXCavBP 1 μM, or TAT-NCXCavBP 10 μM. Kruskal–Wallis test with Dunn's multiple comparison test.

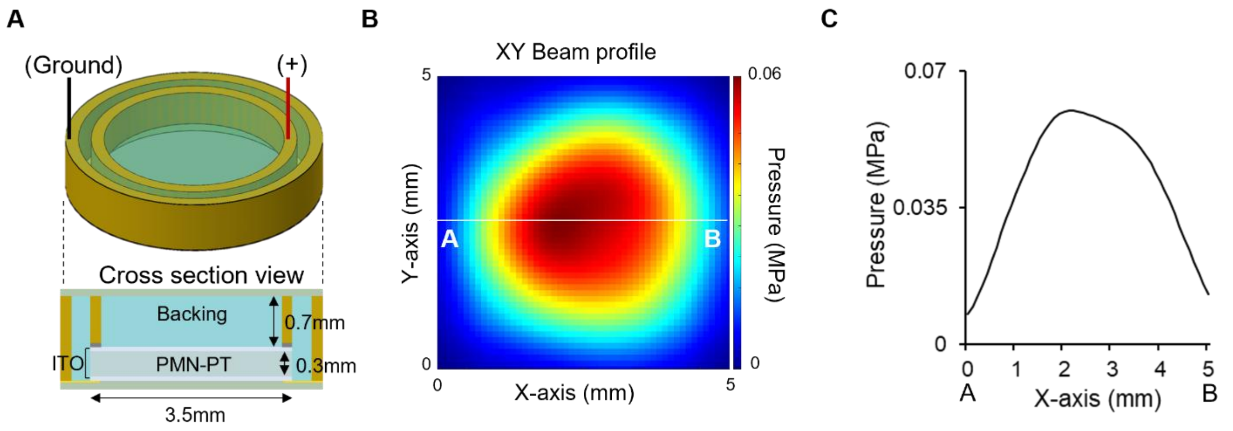


**Supplementary Fig. 7 Acoustic structure of the transparent ultrasonic transducer and its acoustic characterization. a**, Cross-section diagram of the transparent piezoelectric transducer used for in vivo $Ca^{2+}$ imaging experiments. The transducer consists of an ITO transparent electrode layer, a 0.3 mm PMN-PT piezoelectric element, and a 0.7 mm acoustic backing layer, with a total aperture diameter of 3.5 mm. **b**, XY beam pressure profile measured by scanning hydrophone at the focal plane, displayed as a 2D color map (pressure range 0–0.06 MPa). **c**, Acoustic pressure profile (MPa) measured along the A–B horizontal line from (b). The profile demonstrates a Gaussian-shaped lateral beam distribution with a peak pressure of 0.052 MPa.

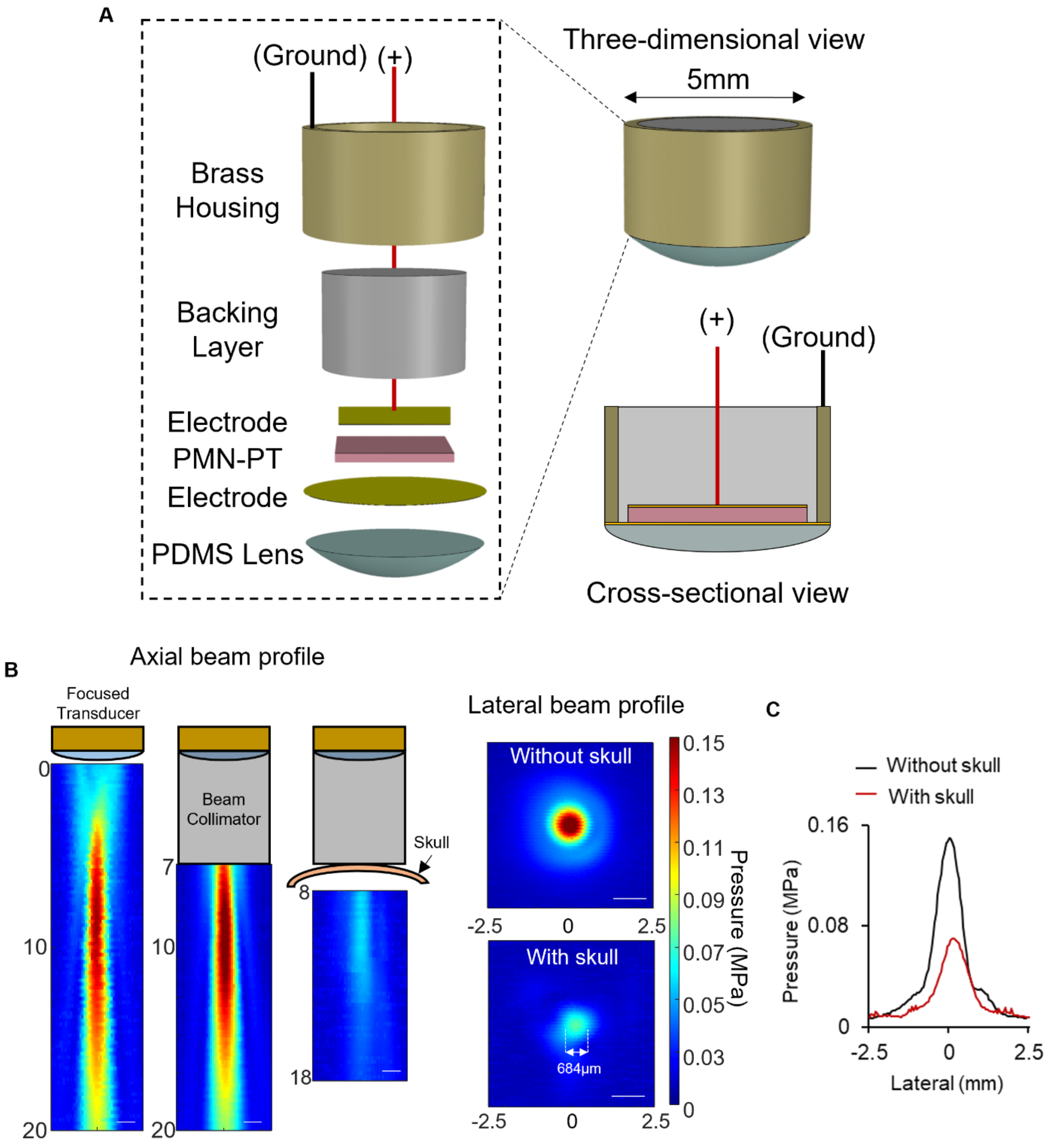


**Supplementary Fig. 8 Structure and acoustic validation of the miniaturized ultrasonic transducer for freely moving mice. a**, Exploded-view schematic (left) and three-dimensional assembled view (right) of the miniaturized wearable focused ultrasound transducer used for in vivo experiments in freely moving mice. Layered components from top to bottom: brass housing, backing layer, electrode, PMN-PT piezoelectric element (pink), electrode, and PDMS acoustic lens. **b**, Axial beam profiles measured for the focused transducer alone, with a beam collimator, and with the collimator plus a mouse skull. Lateral beam profiles (color maps, 0–0.15 MPa) without (top) and with (bottom) skull are shown. Scale bar, 1 mm. **c**, Quantitative lateral pressure profiles at the focus comparing conditions without (black) and with (red) skull. The skull causes modest beam broadening and attenuation while maintaining a narrow focal width of 684 μm.

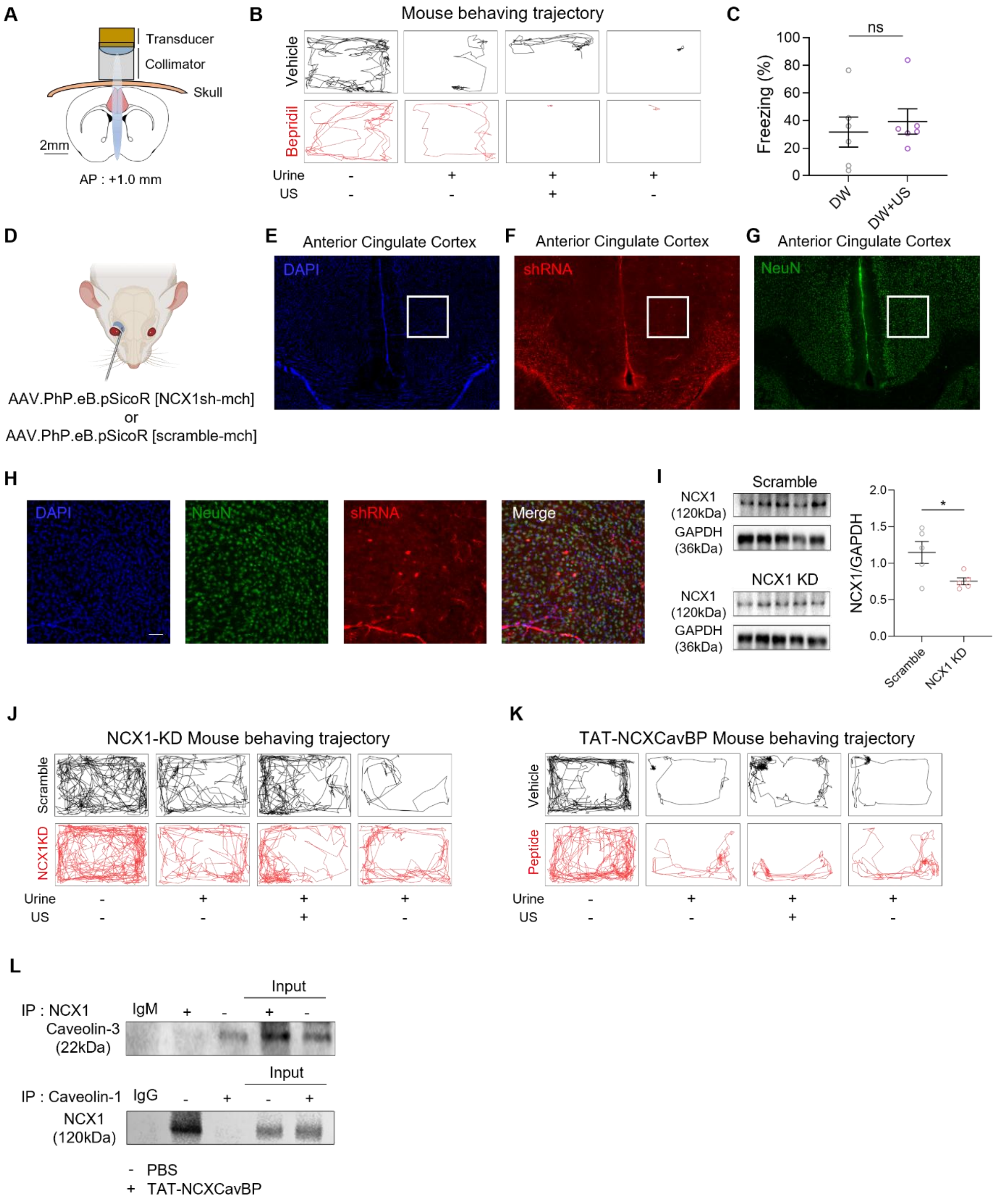


**Supplementary Fig. 9 In vivo validation of viral-mediated NCX1 knockdown, and its functional association with caveolin on behaving trajectories in the ACC. a**, Schematic cross-section showing the miniaturized focused transducer and beam collimator positioned on the intact mouse skull targeting the ACC at AP +1.0 mm from bregma. **b**, Representative movement trajectories of vehicle-treated (black) and bepridil-treated (red) mice across four behavioral conditions. **c**, Freezing percentage comparing distilled water (DW, no odor) and DW+US conditions in naïve mice. Unpaired t-test. n = 6 per group. **d**, Schematic of retro-orbital injection approach. Mice received a retro-orbital injection of AAV.PhP.eB.pSicoR [NCX1sh-mCherry] or AAV.PhP.eB.pSicoR [scramble-mCherry]. **e–g**, Wide-field fluorescence image of the ACC region showing (e) DAPI nuclear staining (blue), (f) shRNA-driven mCherry expression (red) in a coronal brain section, and (g) NeuN immunostaining (green). **h**, High-magnification four-channel fluorescence images of the ACC region showing DAPI (blue), NeuN (green), shRNA-mCherry (red), and merged channels. **i**, Western blot showing NCX1 (120 kDa) and GAPDH (36 kDa) expression in ACC tissue from Scramble and NCX1 KD mice, confirming significant in vivo knockdown. Unpaired t-test. n = 5 per group. **j**, Representative movement trajectories of Scramble shRNA (black) and NCX1 knockdown (NCX1 KD, red) mice across four behavioral conditions: Urine−/US−, Urine+/US−, Urine+/US+, and Urine+/US− (post-stimulation). NCX1 KD mice show impaired ultrasound (US)-mediated reduction in freezing behavior. **k**, Representative movement trajectories of vehicle-treated (black) and TAT-NCXCavBP peptide-treated (red) mice across four behavioral conditions. **l**, Co-immunoprecipitation (IP) analysis of NCX1 interactions with caveolin-3 (upper) and caveolin-1 (lower) in ACC tissue following PBS (vehicle) or TAT-NCXCavBP treatment. IgM and IgG served as negative controls, respectively. TAT-NCXCavBP markedly reduced the association of NCX1 with both caveolin proteins.

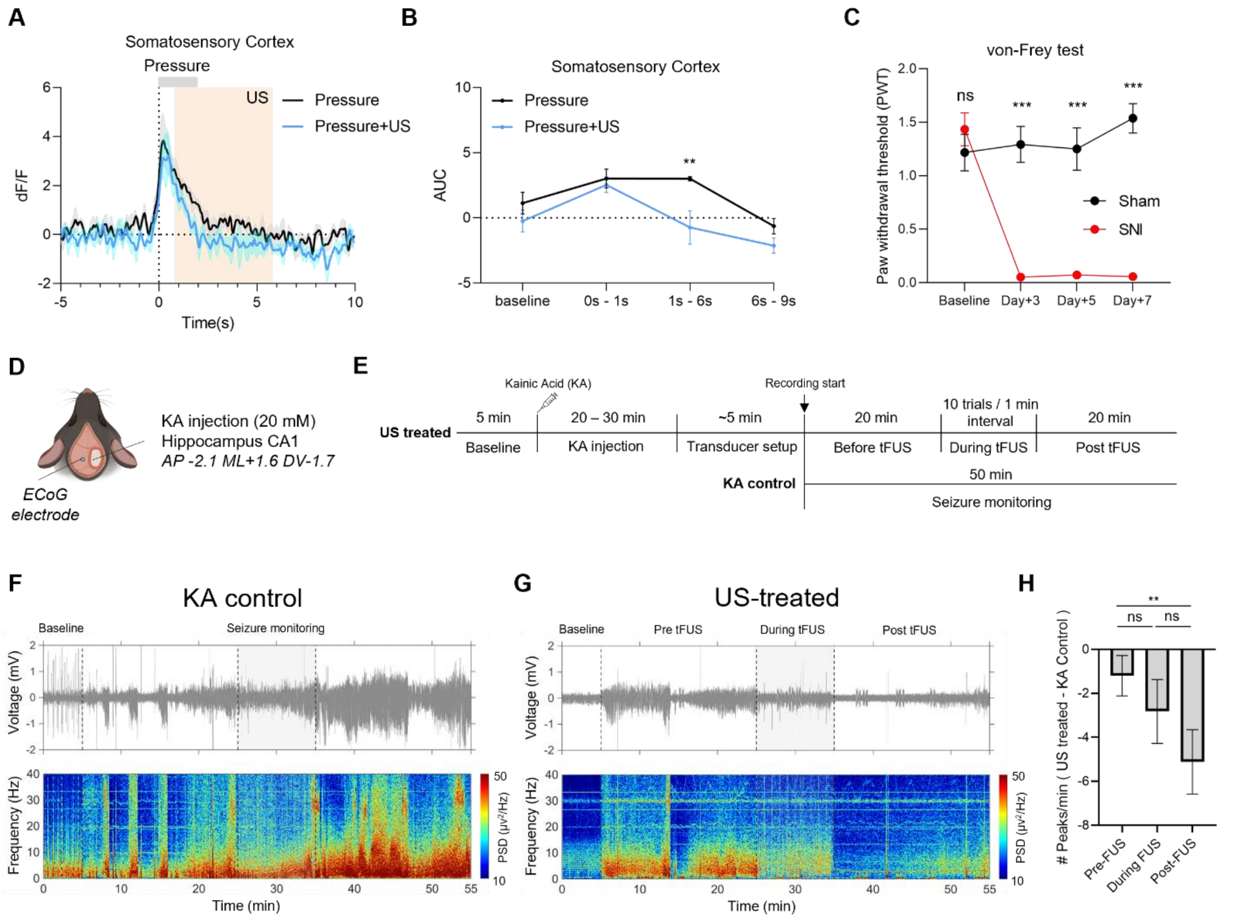


**Supplementary Fig. 10 Fiber photometry recordings of neural activity in the ACC and somatosensory cortex during pressure stimulation and prolonged ultrasound (US) stimulation. a**, Fiber photometry $Ca^{2+}$ traces (dF/F) in the somatosensory cortex comparing pressure alone (black) and pressure+US (blue) in SNI mice. Pressure onset at 0 s; the 5-s US window is shaded. n = 4. **b**, Area under the curve (AUC) analysis of GCaMP6f responses in the somatosensory cortex across four time windows (baseline, 0–1 s, 1–6 s, and 6–9 s). Two-way ANOVA with Sidak's multiple comparison test. n = 4. **c**, Validation of the SNI model by von-Frey testing. Paw withdrawal thresholds in Sham (black, n = 6) and SNI (red, n = 12) mice across post-injury days. Two-way ANOVA with Sidak's multiple comparison test. **d**, Schematic of intrahippocampal kainic acid injection (20 mM, 100 nl) and ECoG electrode placement. **e**, Experimental timeline for the US-treated and KA control group with 50 min of continuous seizure monitoring, including baseline, pre-tFUS, during tFUS (10 trials at 1 min intervals), and post-tFUS. **f**, Representative ECoG traces (top) and corresponding power spectral density (PSD) spectrograms (bottom) from the KA control group. **g**, Representative ECoG trace (top) and PSD spectrogram (bottom) from the US-treated group. **h**, Quantification of the change in seizure peak frequency (US-treated minus KA control) across pre-tFUS, during tFUS, and post-tFUS phases. n = 9. One-way ANOVA with Tukey’s multiple comparison.

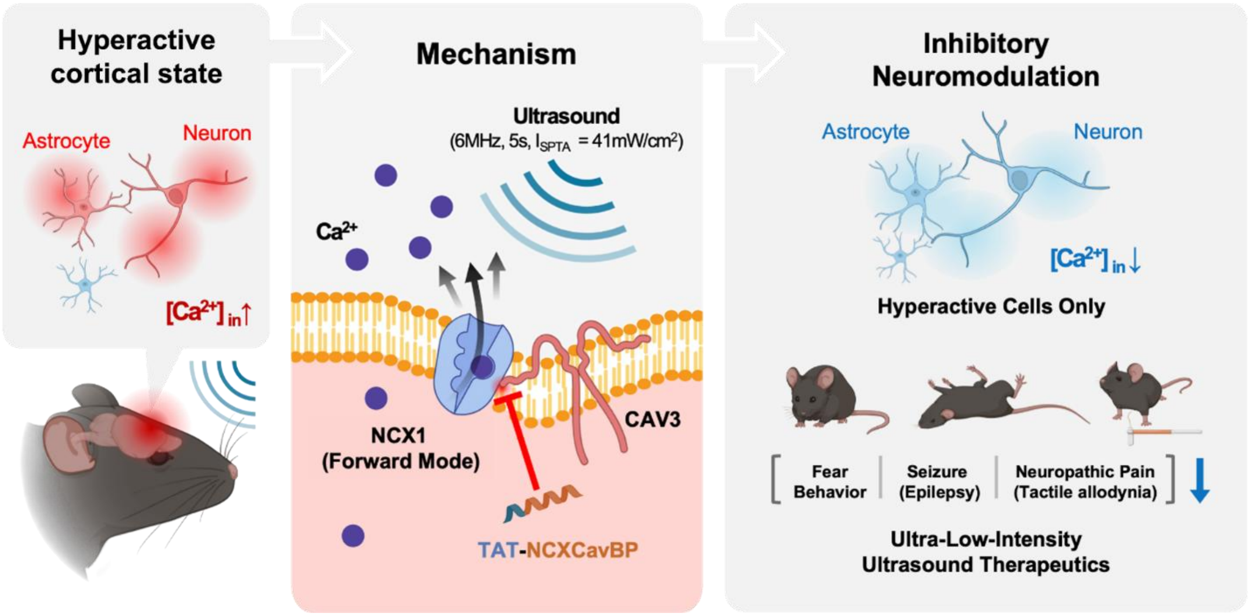


**Supplementary Fig. 11 Graphical summary of the CAV3–NCX1 mechanotransduction axis underlying ultrasound-driven cortical inhibition.** Schematic overview of the proposed mechanism by which prolonged ultra-low-intensity ultrasound stimulation suppresses cortical hyperexcitability. (Left) Under pathological conditions, cortical neurons and astrocytes exhibit elevated intracellular $Ca^{2+}$ concentrations, driving hyperexcitability in brain regions. (Center) Ultrasound stimulation (6 MHz, 5 s, $I_{SPTA}$ = 41 mW/cm²) activates the sodium-calcium exchanger-1 (NCX1) in the forward mode at caveolae microdomains, extruding $Ca^{2+}$ across the plasma membrane. Caveolin-3 (CAV3) serves as the direct NCX1 docking partner and mechanosensitive scaffold that couples acoustic energy to $Ca^{2+}$ extrusion. The TAT-conjugated caveolin-binding peptide TAT-NCXCavBP competitively displaces CAV3 from NCX1, thereby abolishing the ultrasound response. (Right) Activation of the CAV3–NCX1 axis selectively reduces the intracellular calcium concentration ($[Ca^{2+}]_{in}$) in hyperactive cells, suppressing fear behavior, neuropathic pain signaling, and seizure activity in vivo. Thus, the CAV3–NCX1 complex represents a non-pharmacological therapeutic target for conditions of cortical hyperexcitability.
